# Polycystin-1 acts as an atypical adhesion GPCR that responds to novel Wnt signaling and mechanical stimuli

**DOI:** 10.1101/2025.05.21.655326

**Authors:** Nikolay P. Gresko, Valeria Padovano, Peder Berg, Albert C.M. Ong, Stefan Somlo, Nicole Scholz, Tobias Langenhan, Michael J. Caplan

## Abstract

Mutations in the genes encoding Polycystin-1 and 2 cause autosomal dominant polycystic kidney disease (ADPKD). These proteins’ cellular functions and the mechanisms through which their absence causes disease are unknown. We find that the behavior of the 11 transmembrane domain polycystin-1 protein resembles that of an adhesion G protein-coupled receptor when activated by Wnt ligands. Exposure to Wnt ligands causes shedding of the polycystin-1 N terminal region, exposing a tethered agonist that activates G protein-dependent functions of the membrane-associated C terminal fragment. Activated polycystin-1 communicates through G_α13_ to activate RhoA, leading in turn to the ROCK-dependent phosphorylation and inactivation of GSK3β. Activated polycystin-1 traffics out of the primary cilium through an arrestin-dependent mechanism, and arrestin involvement is required for signaling to occur. These data elaborate a function for polycystin-1 that is structurally non-canonical and that is directly connected to pathways whose perturbation results in cystic disease.

**Summary:** The polycystin-1 protein can activate a signaling pathway resembling those employed by non-canonical adhesion G protein coupled receptors, which is induced by Wnt9b binding and which leads to leads to inhibition of the GSK3β kinase and removal of polycystin-1 from the primary cilium.

Graphical Abstract PC1 resides at baseline in the primary cilium. PC1 behaves as an aGPCR that responds to Wnt9b and to flow-induced ATP release by departing from the cilium and initiating the inhibition of GSK3β. The schematic diagram presents a model that integrates the findings presented here. Wnt9b binding to the PC1 NTF leads to shedding of the NTF, exposing the tethered agonist to activate the PC1 receptor-mediated signaling cascade, which involves triggering Gα_13_ binding and the subsequent nucleotide exchange and GTP-loading of RhoA. GTP-bound RhoA stimulates the kinase activity of ROCK, which phosphorylates and inhibits GSK3β.

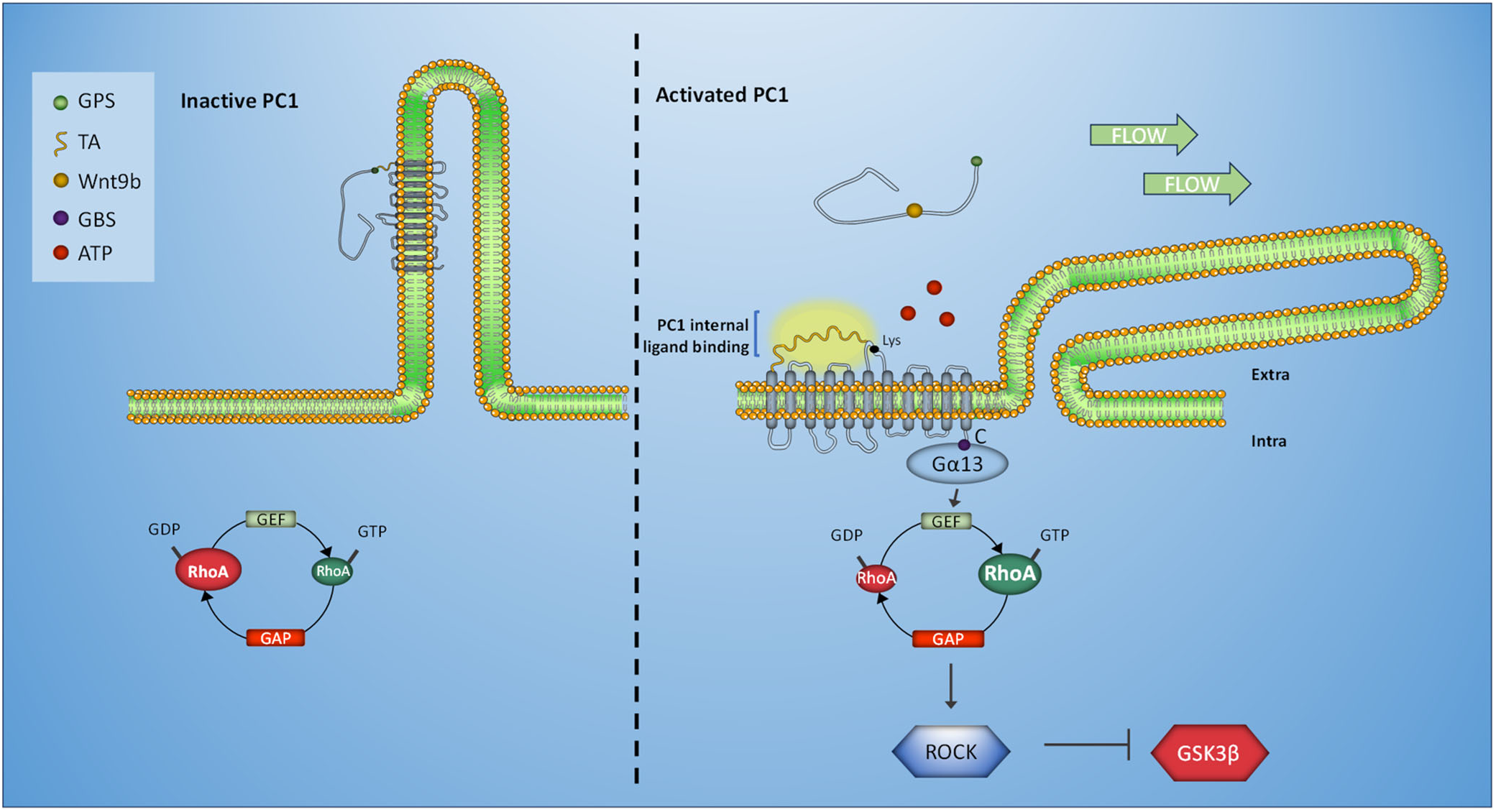

## Introduction

Autosomal dominant polycystic kidney disease (ADPKD) affects ∼1:1,000 people and produces end stage renal disease in 50% of affected individuals. The formation and expansion of nephron-derived fluid-filled cysts compromises the function of the remaining renal parenchyma. Epithelial cells that line cysts are proliferative and mediate active fluid transport into cyst lumens, contributing to cyst expansion^1,2^. Most cases of ADPKD are caused by mutations in *PKD1* or *PKD2*, encoding polycystin-1 (PC1) and polycystin-2 (PC2), respectively. PC1 is a 4,302 amino acid membrane protein that spans the lipid bilayer 11 times. PC2 spans the membrane 6 times and is a member of the transient receptor potential (TRP) family of cation channels^3^. PC1 and PC2 form a heterotetrameric complex^4–6^ that localizes to several subcellular compartments, including the primary cilium^7^ and the mitochondrion-associated membrane domains of the endoplasmic reticulum^8^. Two decades after their discovery, the principal functions of PC1 and PC2 and the processes through which their dysfunction leads to cystic disease remain largely unknown.

The polycystins influence a broad array of signaling cascades, including cAMP, Ca^+2^, G proteins, the mTOR, Wnt, ERK/MEKK pathways and the cell cycle^1,2^. Consequently, their loss leads to perturbations in these and several other processes. It remains unclear which, if any, of these pathways is the primary effector through which polycystins carry out their normal functions. It is clear, however, that dysregulation of many of these pathways contributes to cystogenesis and their activities constitute potential therapeutic targets. One such potential therapeutic target is the GSK3β kinase, which is modulated by numerous signaling pathways, including those responding to insulin and to Wnt ligands^9^. GSK3β activity is elevated in cystic tissue from mouse models of ADPKD, and inhibition of GSK3β slows cyst progression^10,11^.

The short cytoplasmic C terminus of PC1 interacts with trimeric G proteins and deleting the sequence motif that mediates this interaction abrogates critical aspects of PC1 function^12–15^. It has thus been suggested that PC1 is a non-classical G protein coupled receptor (GPCR) for a yet to be identified ligand, despite the divergence of its structure from the canonical GPCR seven transmembrane design^12,13,16^. The possibility that polycystins are receptors also receives support from additional lines of evidence. PC1 undergoes autocatalytic cleavage, producing an N terminal fragment (NTF) that remains non-covalently tethered to the C terminal fragment (CTF), which contains the 11 transmembrane domains and the C-terminal tail^17^. Cleavage occurs within the GAIN (GPCR autoproteolysis-inducing) domain^18^, which is a defining characteristic of adhesion GPCRs (aGPCRs), a large family within the GPCR superfamily^19^. Like PC1, aGPCRs undergo GAIN domain cleavages that generate their large N terminal extracellular regions, which remain non-covalently associated with their transmembrane C terminal components^20,21^. The neo-N terminus of the aGPCR CTF, which is liberated once the NTF is released is referred to as the “*Stachel*”, or synonymously “tethered agonist” (TA), and triggers aGPCR signaling^22,23^. In its inactive conformation the N terminal portion of the GAIN domain envelops the TA, preventing it from interactions with residues within the TM CTF module that activate the receptor. NTF engagement with a stimulatory ligand and/or mechanical stimulation alters its conformation^24^ or causes it to be shed^25,26^, revealing the TA and inducing metabotropic signaling^27^. It has been suggested that PC1 is a non-canonical and orphan member of the aGPCR collection whose ligand and signaling effectors have yet to be identified^28–31^. A mechanism for binding of the putative polycystin-1 TA to its transmembrane domains has been proposed and modeled^31^ ^32^. Finally, the PC1 NTF binds with high affinity to Wnt ligands, including Wnt9b and Wnt5a^33^. These observations suggest that Wnt ligand binding to the PC1 N terminal extracellular region could initiate a conformational transformation that exposes the PC1 TA-equivalent and activates its G protein signaling potential.

We demonstrate that Wnt ligands do indeed unlock the capacity of PC1 to behave like an aGPCR. We show that PC1 signals through the small GTPase RhoA to activate ROCK kinase, which phosphorylates and inhibits GSK3β. Activation of its aGPCR-like function induces the β-arrestin and PKA-dependent removal of PC1 from the primary cilium. These data define a new function for PC1 that connects its intrinsic biological activity directly to a signaling pathway whose perturbation is critically implicated in disease pathogenesis.

## Results

### Wnt9b promotes dissociation of the PC1 NTF from the CTF

Wnt9b binds to the PC1 NTF with an affinity in the low nanomolar range^33^. We hypothesized that, analogous to the behavior of aGPCRs ^21,25,26,28^, this binding will induce dissociation of the PC1 NTF from the CTF. To test this possibility, we stably expressed PC1 and PC2 in HEK293 cells (HEK1+2) and treated them with 13 nM recombinant Wnt9b^33^ for various time intervals. The PC1 construct incorporates a FLAG epitope tag at its N terminus and an HA tag at its C terminus^34^ (Fig. 1a). Blotting with anti-FLAG and anti-HA antibodies reveals that the PC1 protein undergoes very efficient autocatalytic GAIN cleavage, as evidenced by the exceedingly small quantity of full length PC1 (with MW>300 kDa) that is detected in cell lysates by western blotting with the anti-HA antibody (Supp. Fig. 1a). Thus, the majority of the PC1 protein can undergo separation of the NTF from the CTF. Indeed, incubation for 24 hours with Wnt9b reduced the level of the NTF detected in cell lysates significantly without substantially altering the quantity of PC1 CTF (Fig. 1b and Supp. Fig. 1a). Incubation with Wnt9b for 3 hours also reduced significantly, but to a lesser extent, the quantity of the NTF detectable at the cell surfaces, as assessed by anti-FLAG surface immunofluorescence (Fig 1c). Cell surface biotinylation followed by anti-FLAG staining of western blots of proteins recovered in a streptavidin pull down demonstrated that incubation with Wnt9b for as little as 30 minutes led to loss of Flag-tagged PC1 NTF from cell surfaces (Fig. 1d). We have previously shown that incubation in alkaline medium breaks the non-covalent bond that links the NTF to the CTF^34^. As expected, this exposure to “alkaline stripping” similarly reduces the quantity of biotinylated NTF recovered by streptavidin pull down (Fig. 1d). Thus, these data collectively show that Wnt9b promotes physical separation of the PC1 NTF from the CTF.

**Figure 1.**
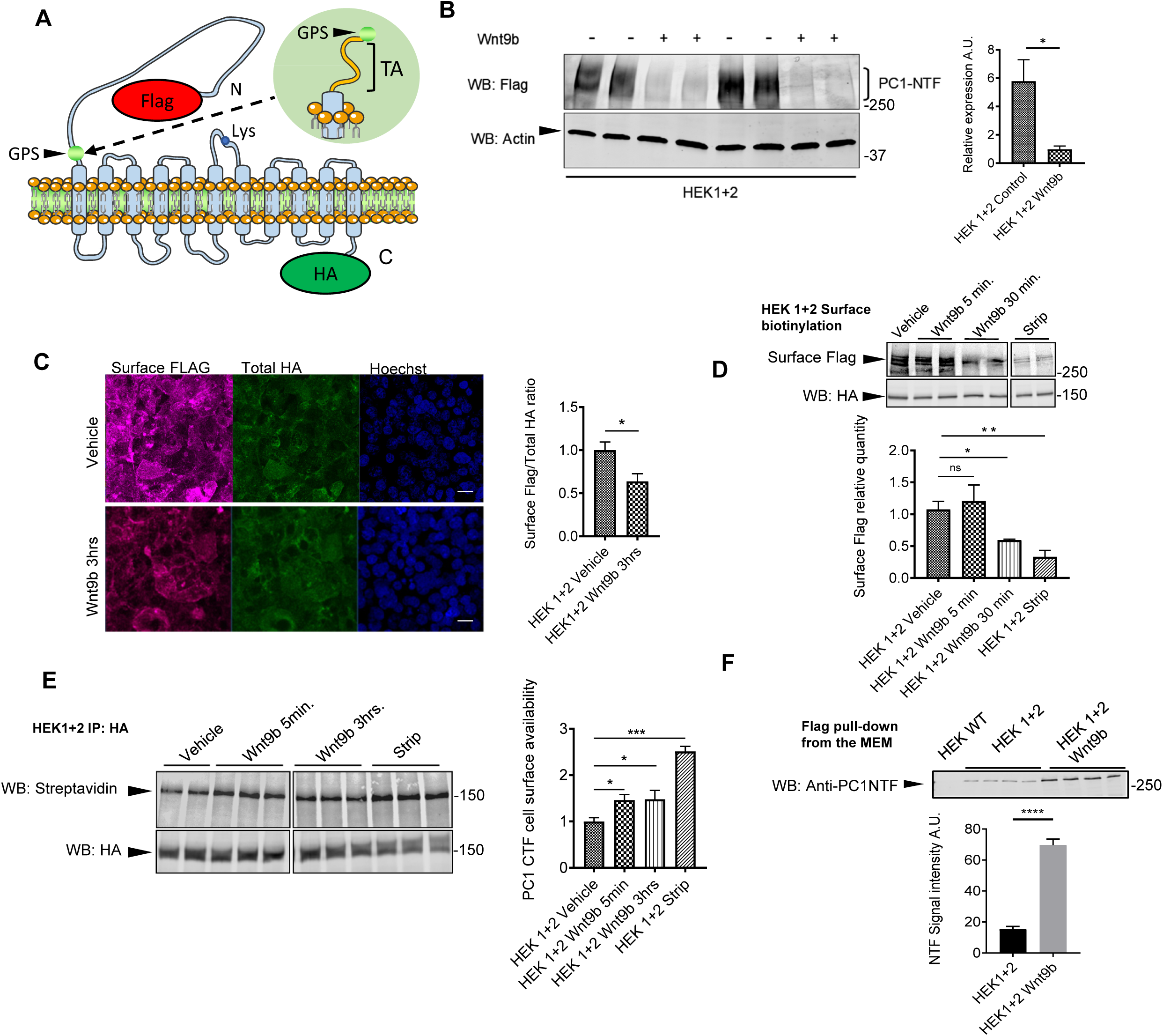
Wnt9b ligand or alkaline pH promote loss of the PC1 NTF from the cell surface. (a) Schematic diagram of the full length PC1 construct employed in these studies. The positions of the Flag tag, HA tag, GPS site and tethered agonist (TA) are indicated, as is the position of the lysine residues whose availability for biotinylation is assessed in 1e. (b) Immunofluorescence detection and quantification of surface PC1 NTF following 3 hrs treatment with Wnt9b. Flag staining (NTF) was performed under non-permeabilizing conditions. HA staining, following permeabilization, revealed total cell quantity of CTF and was used to normalize surface staining. Scale bar=5μM. Graph depicts ratio of Flag to HA staining, normalized to the mean of this value obtained after vehicle treatment. (c) PC1 and 2 expressing-HEK293 cells were treated with Wnt9b or subjected to an alkaline stripping protocol and surface biotinylation was performed. Proteins recovered by streptavidin pull-down were probed with anti-Flag. Cell lysates were probed with anti-HA antibodies. Surface NTF is detected by anti-Flag antibody, and total cell-associated PC1 is represented by the HA signal. The graph depicts ratio of Flag to HA signal, normalized to the mean of this value obtained after vehicle treatment. (d) Western blot detection of total PC1 NTF using anti-Flag antibody. Cells were treated with vehicle or Wnt9b overnight. Bar graphs represent Flag/HA signal ratio. (e) Western blot detection of biotinylated PC1 CTF in experiment where cell surface biotinylation was followed by HA pulldown and streptavidin blotting. Results show increased accessibility of the Lys residue in the PC1-CTF TM6-7 extracellular loop (extracellular loop 3) upon Wnt9b treatment or following alkaline stripping. Total cell-associated PC1 is represented by the HA signal. The graph depicts ratio of Streptavidin to HA signal, normalized to the mean of this value obtained after vehicle treatment. Data are shown as mean ±SEM, n≥3 for all experiments. To assess statistical significance unpaired Student t-Test analysis was performed for all of the panels except for the time course depicted in 1b, for which a paired Student t-Test was employed. P values <0.05 were considered significant. *=P<0.05; **=P<0.01; ***=P<0.001 Differences between mean values obtained for experimental conditions versus control conditions that produce minimal and/or maximal signals were evaluated.

The structure of the PC1 CTF^4^ reveals only one lysine residue that is available to membrane-impermeable Sulfo-NHS-biotin labeling (Fig. 1a). We speculated that the NTF might conceal this residue and that removing the NTF would substantially increase the efficiency of biotinylation of cell surface resident PC1 CTF. Consistent with our hypothesis, we found that biotinylation of the PC1 NTF fell ∼2-fold (Fig. 1e) whereas CTF biotinylation increased ∼3-fold after alkaline stripping (Fig. 1e). Exposure to Wnt9b for 5 minutes or for 3 hours similarly increased CTF biotinylation ∼1.5-fold. Control experiments demonstrate that no biotinylated PC1 protein is detected when cells were transfected with empty vector (EV), and furthermore that the CTF is recovered by streptavidin pull down from cells that express PC1 and 2 but not from cells transfected with EV (Supp. Fig. 1b). Thus, the protein bands detected in these experiments correspond to the PC1 NTF and CTF. As is the case for alkaline stripping^34^, Wnt9b treatment also substantially increased the quantity of the NTF that could be recovered by immunoprecipitation from the medium bathing the cells (Fig. 1f). These observations together demonstrate that Wnt9b promotes dissociation of the PC1 NTF from the CTF, potentially permitting the TA to induce PC1-mediated metabotropic signaling.

### PC1 binds to and inhibits GSK3β

Excessive GSK3β activity has been detected in murine models of ADPKD and in renal cyst tissue from human ADPKD patients, and it has been implicated as a driver of cyst formation^11^. Therefore, we wondered whether PC1 might suppress GSK3β activity, directly or indirectly, in response to Wnt9b binding. We used anti-HA decorated agarose beads to immunoprecipitate PC1 from HEK1+2 cells that had been incubated or not with Wnt9b for various intervals. Western blotting using anti-GSK3β revealed that interaction between PC1 and endogenously expressed GSK3β (Fig. 2a) was profoundly enhanced by incubation with Wnt9b for 3 hours and that this effect occurred in a time-dependent fashion. GSK3β co-precipitation is observed within 5 minutes of the addition of Wnt9b (Fig. 2a, upper panel), a time point associated with displacement of the NTF as revealed by the biotinylation assay depicted in Fig. 1e. Interestingly, a similar association between PC1 and GSK3β is observed when the PC1 NTF was removed by exposure to high pH medium (Fig. 2a, lower panel), suggesting that NTF removal, either by Wnt9b interaction or by alkaline stripping, causes PC1 to associate with a cell signaling protein. Importantly, virtually no GSK3β is detected in precipitates from cells that do not express PC1 (Supp. Fig. 2a). Finally, we also took advantage of the Tg248 *Pkd1* BAC-transgenic mouse line, in which expression of PC1 carrying an N terminal FLAG tag and a C terminal HA tag is under the control of the native *Pkd1* promoter and gene structure. These animals carry 3 copies of the BAC transgenes and likely express the tagged protein at a level that is ∼2-3 fold higher than that of the native PC1^35,36^. Lysates from 8-10 week old BAC-transgenic and wild type mice were subjected to anti-HA immunoprecipitation and the recovered material was analysed by western blotting with anti-GSK3β. While a low level of GSK3β protein is recovered through non-specific adhesion of this protein to the anti-HA beads in the wild type lysate samples, it is clear that substantially more GSK3β is present in the anti-HA immunoprecipitation from lysates prepared from the BAC-transgenic mouse kidneys (Fig. 2b and Supp. Fig. 2b). This experiment demonstrates that PC1 associates with endogenously expressed GSK3β in renal epithelial cells *in vivo*.

**Figure 2.**
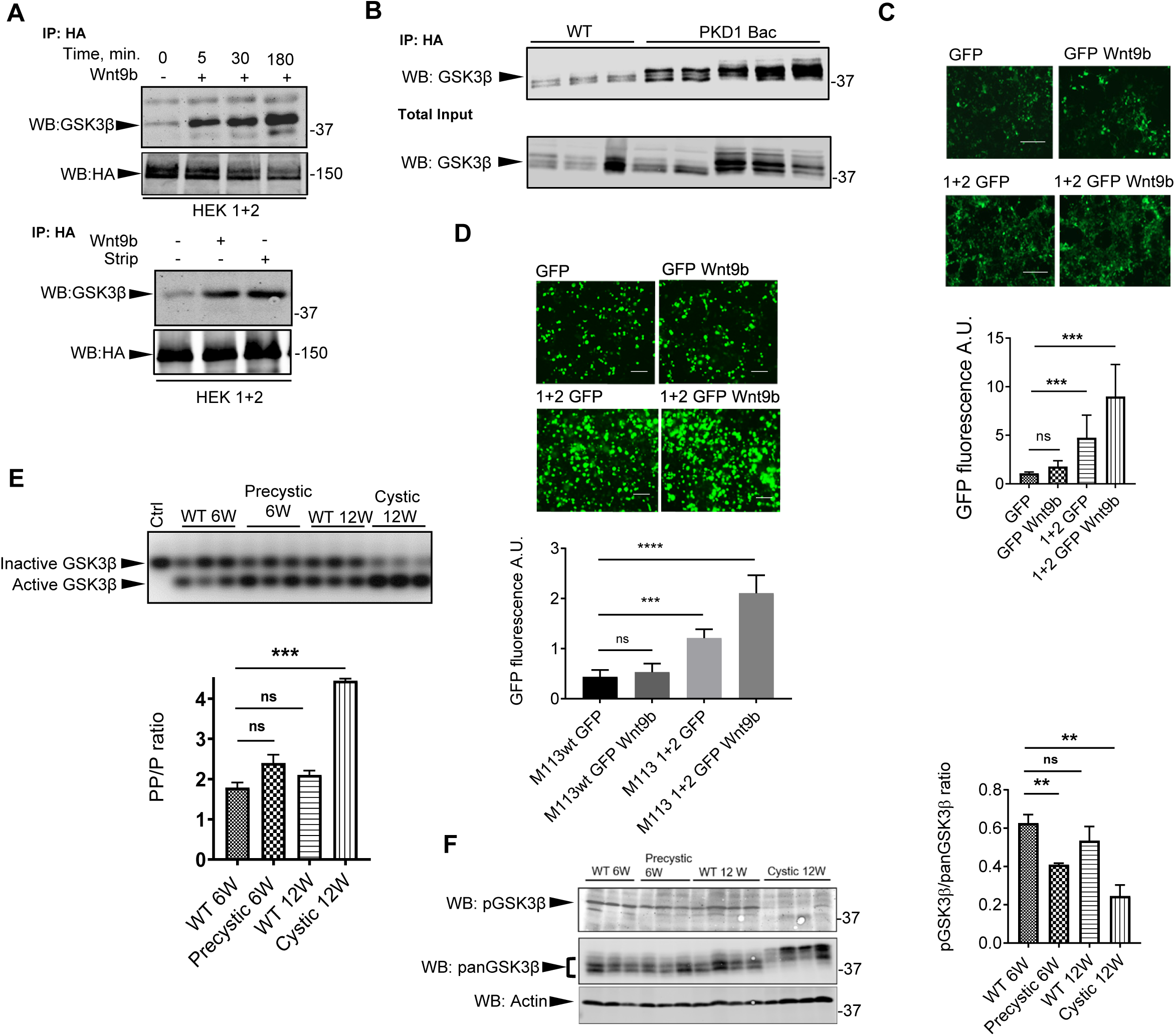
PC1 associates with GSK3β and inhibits its kinase activity. (a) GSK3β co-immunoprecipitates with PC1 following NTF removal by exposure to high pH (strip) or Wnt9b. GSK3β associated with PC1 after 5 minutes of Wnt9b treatment. (b) GSK3β co-immunoprecipitates with PC1 from lysates prepared from Tg248 Pkd1 BAC-transgenic mouse kidneys. (c) Fluorescent biosensor assay of endogenous GSK3β kinase activity in HEK293 cells. PC1 and 2 inhibit GSK3β and this effect is enhanced by 3 hrs treatment with 13nM Wnt9b. Scale bar=100 μM. Graph depicts fluorescence intensity measured in each condition normalized to mean of that measured in cells that express GFP alone. (d) GSK3β kinase assay employing a peptide that electrophoretically migrates in agarose faster when phosphorylated. Lysates from PC1-expressing mouse kidney were compared to similar aged WT kidney lysates. Graph shows mean of the ratio of GSK3β-phosphorylated peptide (PP) to unphosphorylated peptide (P) under each condition. (e) Detection of S9 phosphorylated GSK3β in lysates from un-induced, pre-cystic and cystic kidneys. Blots were probed with anti pan-GSK3β, anti pS9-GSK3β and anti-actin antibodies. Graph depicts the mean of the ratio of the pS9-GSK3β signal to the pan GSK3β signal for each condition. (f) Fluorescent biosensor assay of endogenous GSK3β kinase activity in immortalized murine M113 cells. Untransfected cells were compared to cells expressing PC1 and PC2 as well as to PC1 and 2 expressing cells treated with Wnt9b for 3hrs Scale bar=100 μM. Graph depicts fluorescence intensity measured in each condition normalized to mean of that measured in cells that express GFP alone. Data are shown as mean ±SEM, n≥3 for all experiments. To assess statistical significance unpaired Student t-Test analysis was performed. P values <0.05 were considered significant. *=P<0.05; **=P<0.01; ***=P<0.001 Differences between mean values obtained for experimental conditions versus control conditions that produce minimal and/or maximal signals were evaluated.

To assess PC1 and Wnt9b effects on GSK3β activity in intact cells, we used a fluorescent chimeric biosensor that includes GFP appended to the sequence of a GSK3β substrate^37^. Phosphorylation targets this biosensor for ubiquitination and degradation. Thus, fluorescence intensity is inversely related to endogenous GSK3β activity. Expression of PC1 and 2 significantly reduced endogenous GSK3β activity as indicated by a ∼2-fold increase in GFP fluorescence (Fig. 2c). In cells expressing PC1 and 2, incubation with Wnt9b for 3 hours further inhibited GSK3β by another factor of ∼2. A more complete time course of the GFP signal detected following Wnt9b treatment is presented in Supp. Fig. 2c. Wnt9b produced only a very modest, non-statistically significant effect in mock transfected cells that do not express PC1 and 2, indicative of the specificity of the effect (Fig. 2c). Similarly, Wnt9b treatment did not increase GFP fluorescence when the biosensor was expressed in M113 cells, a *Pkd1*^-/-^ renal epithelial cell line^36^, that is derived from the kidney of a well characterized conditional mouse model of ADPKD (*Pkd1^fl/fl^, Pax8rtTA, Tet-o-Cre*)^38^ (Fig. 2d). In contrast, in M113 cells transfected to express PC1 and PC2, Wnt9b substantially increased GFP fluorescence. These data demonstrate that Wnt9b-induced suppression of GSK3β activity requires polycystin expression in cells derived from an orthologous mouse model of ADPKD.

To determine whether PC1 similarly suppresses GSK3β activity in mouse kidneys *in vivo* we assessed phosphorylation-induced changes in the electrophoretic mobility of a synthetic GSK3β substrate peptide^39^ added to tissue lysates. We prepared lysates from kidneys of a well characterized mouse model (*Pkd1^fl/fl^, Pax8rtTA, Tet-o-Cre*) in which PC1 inactivation can be induced^38^. Six weeks following tet-induction of PC1 inactivation these mice manifest minimal microscopic cystic changes, whereas after twelve-weeks macroscopic cystic disease develops. Because endogenous GSK3β activity is too low to be reliably detected in this assay, we added exogenous GSK3β to the lysates and assessed whether the lysates exert an influence on the activity of this added enzyme. Exogenous GSK3β activity is modestly but significantly greater in lysates from mice six weeks after PC1 inactivation as compared to lysates prepared from un-induced mice (Fig. 2e). Lysates from cystic mice twelve weeks after PC1 inactivation, however, manifest ∼2.5-fold greater exogenous GSK3β activity than that detected in lysates from the un-induced animals. These data suggest that lysates from PC1-expressing renal tissue inhibit exogenously added GSK3β and this inhibitory effect is dependent upon PC1 expression.

Phosphorylation of GSK3β at Ser9 inhibits its kinase activity^40^. We probed mouse kidney lysates prepared at various intervals following PC1 inactivation by western blotting with antibodies that detect phosphorylated S9 GSK3β (pGSK3β) or the entire population of GSK3β protein, which appears as a complex of several bands indicated by the bracket at the left of the panel (Fig. 2f). The fraction of pS9 GSK3β is reduced by ∼30% in lysates from pre-cystic mice six weeks after PC1 inactivation and by ∼65% in lysates from frankly cystic mice twelve weeks after PC1 inactivation as compared to lysates from un-induced controls. It remains to be determined whether the time course of this reduction in pS9 GSK3β is directly linked to the corresponding level of PC1 expression at each time point or if other changes that occur secondary to loss of PC1 expression and the consequent development of cysts influences the level of pS9 GSK3β in this setting. Interestingly, GSK3β migrates with a higher apparent molecular weight in lysates from the cystic mice. The modification responsible for this shift has yet to be elucidated. We also employed immortalized mouse renal epithelial cells that are heterozygous for PC1 expression (Het) or that are homozygous for the deletion of PC1 expression (Null)^41^ (Supp. Fig. 2d). We found by immunofluorescence (Supp. Fig. 2e) and by western blotting (Supp. Fig. 2f) that Het cells manifest increased levels of pS9 GSK3β as compared to PC1 Null cells. Stable re-expression of PC1 in Null cells returns the level of pS9 GSK3β to that detected in Het cells (Supp. Fig. 2f). These data suggest that expression of PC1 in mouse renal epithelial cells *in vivo* and *in vitro* regulates a kinase that mediates an activity-inhibiting phosphorylation of GSK3β on Ser9.

### Truncated PC1 lacking the NTF is constitutively active

By analogy with aGPCRs^20,22,23,28,42–44^ we hypothesized that PC1 protein lacking its NTF would constitutively activate its downstream signaling pathways. We generated truncated PC1 lacking its NTF (PC1^ΔNTF^). The design of the construct includes an N terminal HA sequence, followed by a sequence from the P2Y12 receptor to facilitate its proper targeting, followed by the sequence of the PC1 CTF beginning with the Thr-Ala-Phe (TAF) residues, with the Thr residue corresponding to the +1 residue of the GPS. This design has been used and extensively validated in studies of aGPCRs^22,23^. While this protein does not contain an N terminal signal peptide, this is the case for many multispanning membrane proteins including the vast majority of GPCRs (including aGPCRs), for which signal anchor sequences ensure correct protein targeting to the secretory pathway and membrane topogenesis^45,46^. Schematic diagrams of the design of PC1^ΔNTF^ and its related protein constructs is presented in Supp. Fig. 3a. When transiently overexpressed in HEK293 cells without any added PC2, PC1^ΔNTF^ strongly co-precipitates with GSK3β, much like wild type PC1 that has been activated by Wnt9b incubation (Fig. 3a). Whereas the full length PC1 protein requires co-expression of PC2 to traffic to the plasma membrane^47^, both the PC1^ΔNTF^ and the PC1^ΔNTF+ΔTA^ proteins are delivered to the cell surface without need for co-expression of PC2 as revealed by a surface anti-Flag immunofluorescence labelling assay performed on non-permeabilized cells (Fig. 3b). Previous studies demonstrate that essentially no full length PC1 is delivered to the cell surface in the absence of co-expression with PC2^47–49^.The fluorescent GSK3β activity assay demonstrated that PC1^ΔNTF^ significantly reduces the activity of endogenous GSK3β to a level equivalent to that achieved through incubation with a saturating concentration of the GSK3β inhibitor SB216763 (Fig. 3c). Expression at similar levels of a version of the PC1^ΔNTF^ protein that lacks the tethered agonist (PC1^ΔNTF+ΔTA^) does not inhibit the kinase activity of GSK3β (Fig. 3c, and Supp. Figs. 3a and b), suggesting a TA-dependent activation mechanism. Similarly, significantly less GSK3β protein co-precipitates with PC1^ΔNTF+ΔTA^ as compared to that which associates with the constitutively active PC1^ΔNTF^ (Fig. 3a).

**Figure 3.**
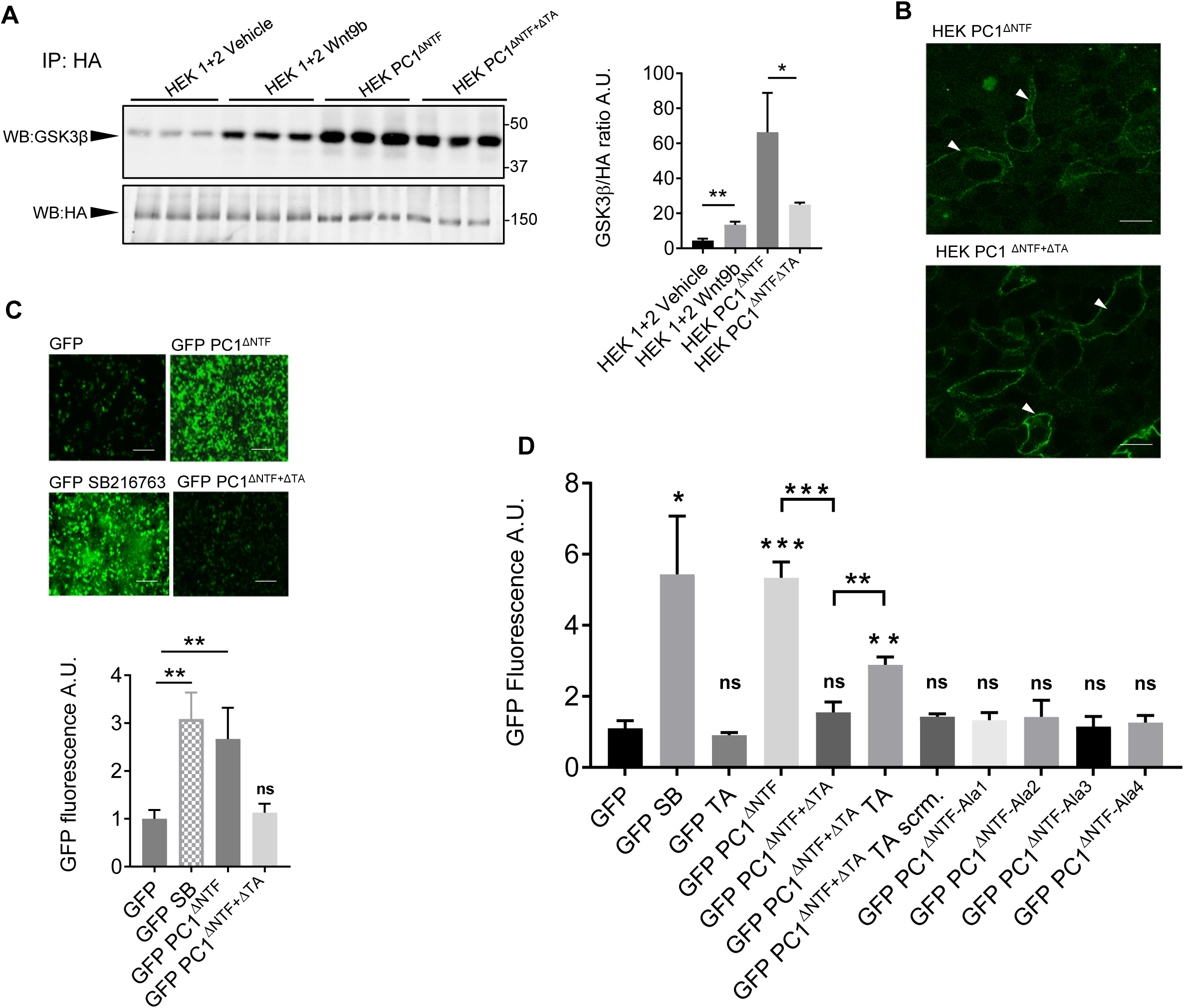
PC1 lacking its NTF is constitutively active. (a) PC1^ΔNTF^ co-immunoprecipitates with GSK3β to the same extent as Wnt9b-stimulated full length PC1. Less GSK3β co-immunoprecipitates with PC1^ΔNTF+ΔTA^, which lacks a tethered agonist. The graph depicts the ratio of GSK3β signal to the HA signal, normalized to the value of this ratio for the HEK 1+2 condition. (b) GSK3β biosensor shows that PC1^ΔNTF^ inhibits GSK3β to the same extent as a maximal concentration (10 μM) of GSK3β inhibitor SB-216763. PC1^ΔNTF+ΔTA^ does not inhibit GSK3β. The graph depicts the ratio of GFP fluorescence in each condition, normalized to the mean of this value obtained with cells that express GFP alone. (c) Confocal Z-stack imaging of PC1^ΔNTF^ and PC1^ΔNTF+ΔTA^ detected with anti-HA antibody added to the media bathing unfixed and non-permeabilized cells shows that both proteins expressed in the absence of PC2 reach the surfaces of HEK293 cells. (d) PC1^ΔNTF+ΔTA^, lacking the Stachel sequence, does not inhibit the GSK3β in the fluorescence biosensor assay. The ability of PC1^ΔNTF+ΔTA^ to inhibit GSK3β can be partially restored by addition of soluble Stachel peptide (300 μM). Scrambled Stachel sequence (scrm) did not produce this effect. Versions of the PC1^ΔNTF^ protein that carry successive groups of alanine substitutions in the sequences of their tethered agonists (PC1^ΔNTF^-Ala1-4) do not inhibit GSK3β. Data are shown as mean ±SEM, n≥3 for all experiments. To assess statistical significance unpaired Student t-Test analysis was performed. P values <0.05 were considered significant *=P<0.05; **=P<0.01; ***=P<0.001.; ****=P<0.0001 Differences between mean values obtained for experimental conditions versus control conditions that produce minimal and/or maximal signals were evaluated. In panel 3d the values obtained for each condition were compared to those obtained in the GFP alone condition (indicated by the symbols above each bar) and pairwise comparisons of the values obtained in a subset of conditions related by a single experimental manipulation are indicated by the brackets above the bars.

aGPCRs that lack both their NTF and TA can be activated by high concentrations of soluble Stachel peptides that correspond to the sequence of the element within the native receptor protein TA/Stachel sequence^22,23^. Incubating cells that express PC1^ΔNTF+ΔTA^ in medium containing putative PC1 TA peptide (100 and 300 μM) but not with a scrambled version of this peptide, partially recapitulates the GSK3β inhibition produced by expression of PC1^ΔNTF^ (Fig. 3d and Supp. Fig. 3c). These results are consistent with those recently obtained in studies using an assay system that reports on a different potential PC1 effector signaling pathway^32^. Furthermore, expression of each of 4 versions of PC1^ΔNTF^ protein carrying successive groups of alanine substitutions of 6 adjacent residues of the 24-residue putative TA sequence (PC1^ΔNTF-Ala1–4^) (depicted graphically in Supp. Fig. 3a) does not inhibit GSK3β (Fig. 3d). These data together provide strong support for a model in which PC1 activation resembles a mechanism that operates for an aGPCR. Removal of its NTF, induced by Wnt9b binding or as a consequence of genetic manipulation or of alkaline stripping, exposes a TA and thereby unleashes the capacity of the PC1 protein to suppress GSK3β activity.

### Activated PC1 recruits G_α13_ and engages the RhoA-ROCK pathway to inhibit GSK3β

PC1 interacts with several trimeric G protein α-subunits^12–15^. To identify molecular players involved in PC1-mediated GSK3β inhibition, we used the fluorescent GSK3β activity assay combined with selective inhibition of components of G protein signaling pathways that are employed by aGPCR family members, including G_α12_ and G_α13_, RhoA and the RhoA-activated kinase ROCK^43,50–53^. ROCK phosphorylates GSK3β at Ser9 and thus constitutes a potential link between activation of the PC1 receptor function and inhibition of GSK3β^54^.

In cells that express PC1 and 2 as well as the fluorescent biosensor that also received RhoA-targeted siRNA, Wnt9b did not induce inhibition of GSK3β beyond the level that was seen in PC1 and 2 expressing cells that were not Wnt9b-treated (Fig. 4a and Supp. Fig. 4a). Scrambled siRNA did not produce this suppressive effect. Substantial reductions in GFP fluorescence were also seen when cells were treated with the chemical ROCK inhibitor Y27632 or when they were treated with siRNA directed against G_α13_ (Fig. 4a and Supp. Fig. 4b). The inhibitory effects of PC1^ΔNTF^ expression on GSK3β were suppressed to a remarkably similar extent by RhoA knockdown, ROCK inhibition and G_α13_ knockdown (Fig. 4a).

**Figure 4.**
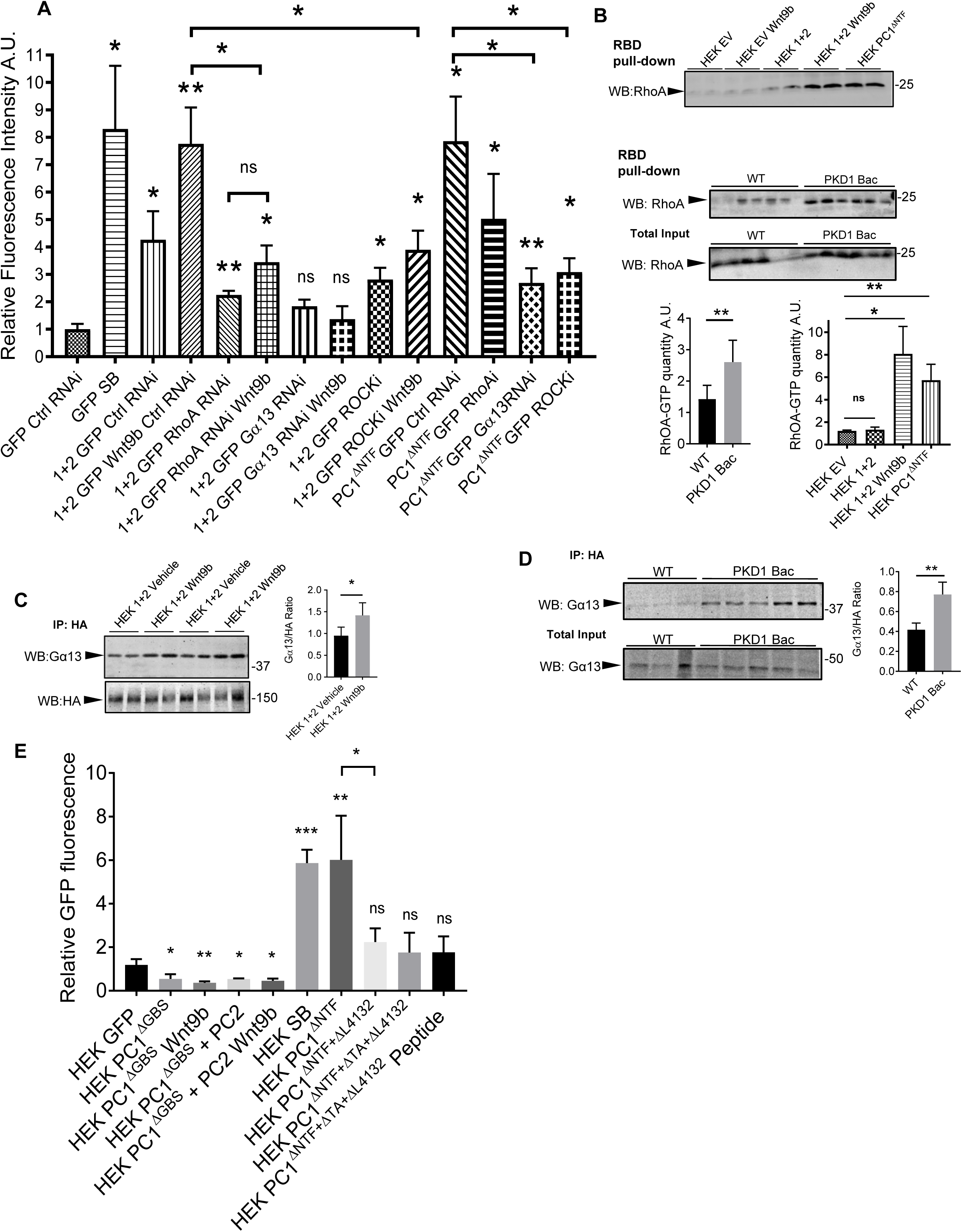
PC1-mediated GSK3β inhibition involves Gα13-RhoA-ROCK. **(a)** GSK3β activity measured +/− Gα13 or RhoA or control (Ctrl) siRNAs, or ROCK inhibitor Y27632. Suppressing Gα13, RhoA or ROCK dampened GSK3β inhibition by Wnt9b-stimulated PC1 or PC1ΔNTF. 10 μM SB-216763 (SB) treatment demonstrates fluorescence detected with maximal GSK3β inhibition. Graph depicts means of ratios of GFP signals normalized to mean of value obtained with cells transfected with GFP and Ctrl RNAi. **(b)** GTP-bound RhoA abundance was assessed by rotekin RhoA-GTP binding domain bead pull-down, followed by western blotting for RhoA. Wnt9b stimulation of PC1, or PC1ΔNTF expression, increased GTP-bound RhoA. Graph depicts means of ratios of RhoA-GTP signal, normalized to mean of value obtained with empty vector (EV) transfection. The quantity of RhoA-GTP is also substantially higher in lysates prepared from the kidneys of Tg248 Pkd1 BAC-transgenic mice as compared to that present in lysates of wild type mouse kidneys. **(c)** and **(d)** Detection of *in vitro* (c) and *in vivo* (d) quantity of Gα13 present in anti-HA precipitates of PC1 was assessed by western blotting. Wnt9b treatment modestly but significantly increased quantity of PC1-associated Gα13 PC1 (c). Graphs in c and d depict means of ratios of signal detected in Wnt9b-treated condition, normalized to mean of value obtained with vehicle-treated cells. Graph depicts the intensity of the Gα13 signal in each condition normalized to the corresponding HA signal. **(e)** Biosensor assay to detect importance of G-protein binding site of PC1 for regulation of GSK3β. PC1 or PC1^ΔNTF^ lacking its G-protein binding site, or PC1 carrying a mutation within the G-protein binding site sequence (ΔL4132), does not inhibit GSK3β. Similarly, soluble Stachel peptide does not induce GSK3β inhibition in cells expressing PC1^ΔNTF+ΔTA+ΔL4132^. Graph depicts ratio of GFP fluorescence, normalized to mean of value obtained with cells expressing GFP alone. Data are shown as mean ±SEM, n≥3 for all experiments. To assess statistical significance unpaired Student t-Test analysis was performed. P values ≤ 0.05 were considered significant *=P<0.05; **=P<0.01; ***=P<0.001.; ****=P<0.0001 Differences between mean values obtained for experimental conditions versus control conditions that produce minimal and/or maximal signals were evaluated. In panels 4a and 4f the values obtained for each condition were compared to those obtained in the GFP alone condition (indicated by the symbols above each bar) and pairwise comparisons of the values obtained in a subset of conditions related by a single experimental manipulation are indicated by the brackets above the bars.

Next we conducted a pull-down assay that employed rhotekin covalently attached to agarose beads^55^. Rhotekin binds tightly to the GTP bound form of RhoA. Lysates from HEK293 cells that did or did not express PC1 and 2 or PC1^ΔNTF^ and that were or were not treated with Wnt9b were subjected to rhotekin agarose pull down and western blots of the recovered material were probed with anti-RhoA. Wnt9b treatment of cells expressing PC1 and 2 led to an ∼8-fold increase in the quantity of GTP-bound RhoA (Fig. 4b, top panel). Interestingly, a similar enhancement in the levels of GTP-bound RhoA was seen in cells that express PC1^ΔNTF^ that had not been exposed to Wnt9b. We also performed this experiment using lysates prepared from Tg248 *Pkd1* BAC transgenic mice, which express ∼3-fold higher levels of PC1 than do wild type animals^35,36^. A similar ∼3-fold enhancement in the fraction of GTP-bound RhoA was seen in lysates from the Tg248 mice as compared to the level detected in lysates prepared from wild type animals (Fig. 4b, middle panel). Finally, we performed a co-immunoprecipitation experiment that showed that PC1 interacts with the α-subunit of G_α13_. (Fig. 4c and d). G_α13_ was not detected in anti-HA immunoprecipitates from cells that were transfected with empty vector (Supp. Fig. 4c). G_α13_ was also detected in anti-HA immunoprecipitates prepared from lysates of kidneys from the Tg248 *Pkd1* BAC transgenic mice, whereas very little G_α13_ was present in anti-HA immunoprecipitates prepared from lysates of wild type kidneys (Fig. 4d). The co-precipitation of G_α13_ with PC1 was significantly (∼1.5 fold) stimulated by Wnt9b treatment of transfected HEK293 cells (Fig. 4c). Like wild type PC1, the PC1 protein lacking its putative G protein binding site (PC1^ΔGBS^)^13^ co-expressed with PC2 reaches the cell surface (Supp. Fig. 4d). Furthermore, PC1^ΔGBS^ does not inhibit GSK3β activity in the presence or absence of Wnt9b (Fig. 4e and Supp. Fig. 4e). A pathogenic single residue deletion mutation (ΔL4132) immediately adjacent to the PC1 protein’s G protein binding motif interferes with G protein binding to PC1^12^. We created versions of PC1^ΔNTF^ and PC1^ΔNTF+ΔTA^ that carry this same deletion. Although PC1^ΔNTF+ΔL4132^ contains the TA, it does not inhibit GSK3β the fluorescence assay (Fig. 4e and Supp. Fig. 4e). Similarly, addition of soluble Stachel TA peptide does not induce PC1^ΔNTF+ΔTA+ΔL4132^ to inhibit GSK3β (Fig. 4e). Thus, PC1 must be capable of interacting with G proteins in order for it to inhibit GSK3β in response to Wnt9b, to removal of its N terminus or to addition of exogenous TA peptide. Furthermore, TA stimulation of PC1 at the extracellular side hierarchically precedes intracellular activation of G_α13_. These data strongly suggest that G_α13_, RhoA and ROCK are obligate participants in the PC1-dependent, Wnt9b-stimulated inhibition of GSK3β.

### PC1 receptor activity governs its localization to cilia

Activated GPCRs are subjected to a number of mechanisms that lead to their desensitization, which can include phosphorylation and β-arrestin-dependent endocytosis^56^. In the case of GPCRs that reside in the primary cilium, ligand binding and receptor activation can induce receptor trafficking out of the cilium through processes that can involve β-arrestin, ubiquitination and components of the BBSome ciliary trafficking complex^57,58^. To assess whether and how the receptor-like properties of PC1 might affect its ciliary localization we made use of the LLC-PK_1_ line of porcine kidney epithelial cells. These cells have previously been used to study the post-synthetic trafficking of PC1 to the primary cilium^34^. When expressed in LLC-PK_1_ cells, the PC1 and 2 proteins co-localize with the ciliary marker ARL13b in the primary cilium. Incubating PC1 and 2-expressing LLC-PK1 cells with Wnt9b for 24 hours results in a substantial reduction in the ciliary pool of PC1 (Fig. 5a). When cells were treated with the β-arrestin inhibitor barbadin^59^ the size of the ciliary PC1 pool was unchanged upon Wnt9b stimulation (Fig. 5a). These observations suggest that Wnt9b-dependent PC1 activation leads to its β-arrestin mediated removal from the primary cilium.

**Figure 5.**
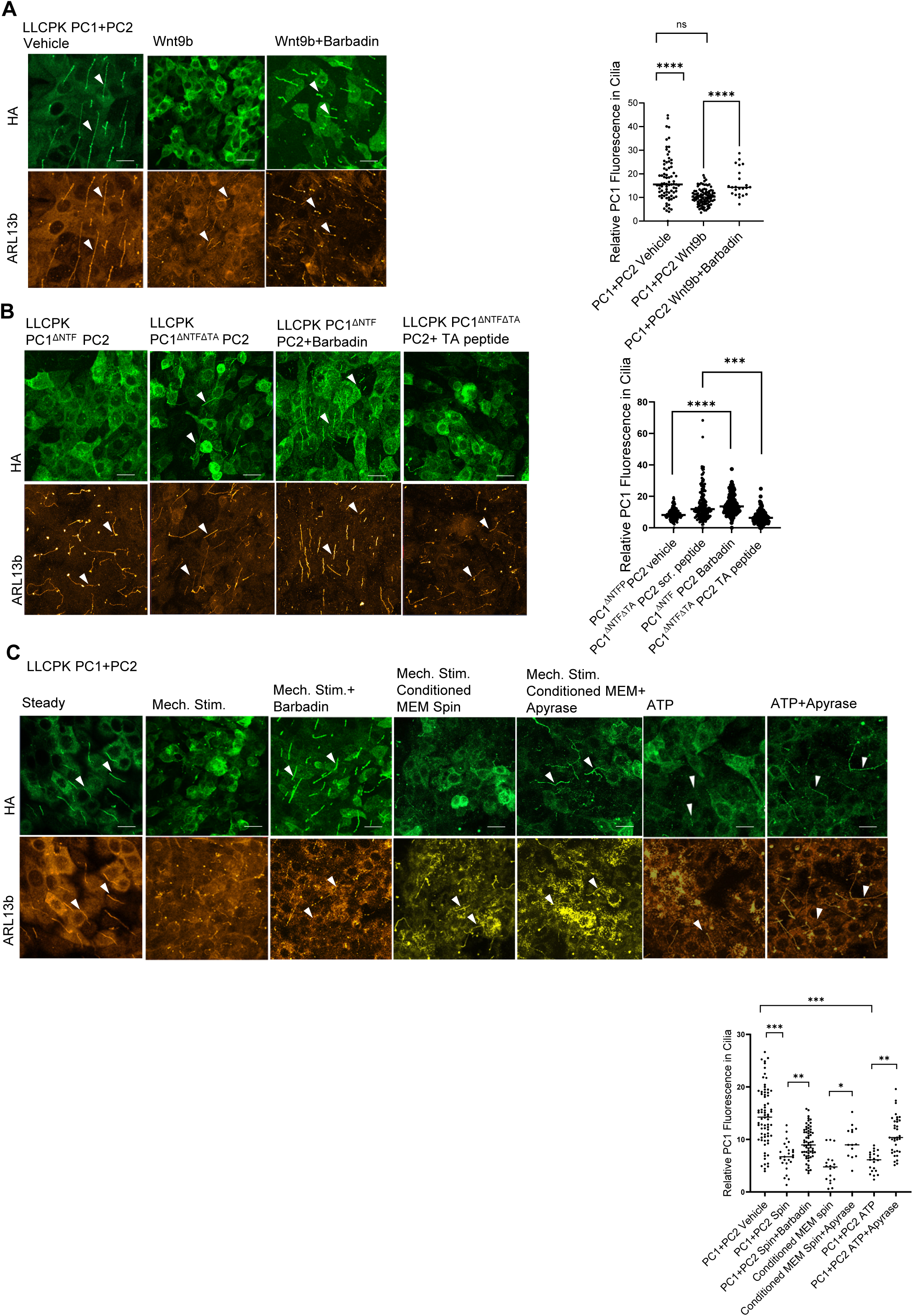
PC1 activity controls its localization to the primary cilia. **(a)** Immunofluorescent localization of the full length PC1 (green) in the primary cilia of LLCPK1+2 cells treated with wnt9b, or with a combination of Wnt9b and barbadin. Cilia are detected by immunostaining for Arl13b (orange) **(b)** Immunofluorescent localization of PC1^ΔNTF^ and PC1^ΔNTFΔTA^ in the primary cilia of LLCPK cells. Barbadin treatment results in localization of PC1^ΔNTF^ to cilia, while treatment with soluble TA peptide causes loss of PC1^ΔNTF+ΔTA^ from the cilia. **(c)** Immunofluorescence detection of the full length PC1 in the primary cilia of cells treated with mechanical stimulation, with medium from cells preconditioned by mechanical stimulation or with ATP added to the medium. Barbadin and Apyrase treatment blocked the effects of mechanical stimulation and ATP on PC1 ciliary localization. Data are shown as mean ±SEM, n≥3 for all experiments. Scale bar = 5μ. To assess statistical significance unpaired Student t-Test analysis was performed. P values <0.05 were considered significant *=P<0.05; **=P<0.01; ***=P<0.001.; ****=P<0.0001 Differences between mean values obtained for experimental conditions versus control conditions that produce minimal and/or maximal signals were evaluated.

The data presented in Fig. 3c and 3d demonstrate that the PC1^ΔNTF^ protein behaves as a constitutively activated PC1 receptor, while PC1^ΔNTFΔTA^ behaves as a constitutively inactive form of the receptor that can be activated through the exposure to a soluble form of the TA. Consistent with this behavior, we find that PC1^ΔNTF^ protein expressed in LLC-PK1 cells is absent from primary cilia (Fig. 5b). When these cells are incubated with barbadin for 18 hours the PC1^ΔNTF^ protein is prominently localized to cilia. Conversely, PC1^ΔNTFΔTA^ localizes to cilia in untreated cells, while incubation with soluble TA peptide for 18 hours leads to a marked reduction in this protein’s ciliary localization (Fig. 5b). Taken together, these data lend further to support to the interpretation that the ciliary localization of PC1 is governed by the status of its activity.

A number of studies suggest that the polycystins may participate in and respond to ciliary mechanosensation^60–68^. Since we find that Wnt ligand activation induces PC1 to depart from the cilium, we wondered whether mechanical stimulation would have a similar effect. LLC-PK1 cells expressing PC1 and PC2 were grown to confluence on glass cover slips placed at the bottom of the wells of 24 well plates. Plates were placed, or not, on an orbital shaker set to 100 rpm and maintained in a 37°C incubator. After 48 hours the cover slips were fixed and processed for immunofluorescence using antibodies directed against the HA epitope to detect PC1 and against the cilia marker ARL13B. As can be seen in Fig. 5c, in cells that were subjected to orbital shaking the quantity of PC1 in cilia is substantially reduced as compared to that detected in control cells that were maintained under static conditions. Addition of barbadin to the medium bathing the cells subjected to shaking prevented the loss of PC1 from cilia (Fig. 5c).

### Effect of ATP on PC1 ciliary localization and signaling

If PC1 were acting as a ciliary mechanosensor that is directly activated by the fluid shear force created by orbital shaking, then we would expect that the effects of shaking on PC1 localization would manifest rapidly. Perhaps surprisingly, the effects of mechanical stimulation on PC1 ciliary localization manifest slowly, becoming detectable only after at least 48 hours of shaking (Supp. Fig 5a). This behavior suggested the possibility that the effects of shear stress on PC1 ciliary localization could be indirect and that they might perhaps be mediated by a factor that is secreted into the medium in response to shear stress. Were this the case, the slow time course that we observed could reflect the need for this factor to accumulate to a sufficient concentration in the relatively large volume of medium bathing the cells. To test this possibility, we prepared medium condition by PC1 and PC2-expressing LLC-PK1 cells grown in 10 cm dishes and subjected or not to 48 hours of orbital shaking (25 rpm). When applied to cells grown under static conditions, medium conditioned for at least 48 hours by shaken cells induced the departure of PC1 from the cilium (Fig. 5c and Supp. Fig. 5a). The effect of the medium conditioned by shaken cells was blocked by barbadin. In addition, medium conditioned by shaken cells that did not express PC1 and PC2 did not alter the ciliary localization of PC1 in the cells to which it was applied (Supp. Fig. 5b). Taken together, these data suggest that mechanical stimulation of polycystin-expressing cells induces the secretion of a factor into the medium. This factor may subsequently trigger the GPCR-like function of PC1, resulting in its β-arrestin-driven removal from the cilium.

Bending the primary cilium has been shown to induce vesicular ATP release in renal epithelial cells in culture and in the intact nephron^69–71^. We wondered, therefore, whether ATP could be the factor that “conditions” the medium from shaken polycystin-expressing LLC-PK1 cells. To test this possibility, we treated the conditioned medium with apyrase to degrade ATP and applied it to unshaken PC1 and PC2-expressing LLC-PK1 cells. The apyrase-treated conditioned medium did not induce PC1 to depart from the cilium. In contrast, adding 1 mM ATP to the medium bathing unshaken polycystin-expressing LLC-PK1 cells resulted in the departure of PC1 from primary cilia within 3 hours of incubation (Fig. 5c). We also assessed the effects of ATP addition on PC1-mediated signaling in the GSK3β inhibition assay. Adding ATP to the medium bathing biosensor-expressing HEK293 cells induces the PC1 and 2-dependent inhibition of GSK3β, with half maximal inhibition observed in the presence of ∼10 µM extracellular ATP. In cells that express PC1^ΔGBS^ (which lacks the G protein binding site) in association with PC2, the addition of ATP to the medium did not induce inhibition of GSK3β (Sup. Figs. 6a and 6b).

The Polycystin-1, Lipoxygenase, Alpha-Toxin (PLAT) domain of PC1 has been shown to interact with β-arrestin and to mediate the β-arrestin-dependent internalization of PC1 from the plasma membrane^72^. This interaction and its effects on PC1 localization are dependent upon the PKA-mediated phosphorylation of serine 3164. To explore the possible role of this phosphorylation-dependent β-arrestin interaction in modulating the ciliary localization of PC1, we expressed non-phosphorylatable S3164A or phosphomimetic S3164D versions^72^ of PC1 together with PC2 in LLC-PK1 cells. The S3164A PC1 protein localized to cilia, and this localization was not influenced by incubation with ATP. In contrast, the S3164D PC1 protein was not detected in cilia under baseline conditions, but incubation with barbadin resulted in its ciliary accumulation (Fig. 6a). Finally, the co-incubation with the PKA inhibitor H89 (5 µM) blocked the departure of wild type PC1 protein from primary cilia in response to ATP treatment (Fig. 6a). Taken together, these data support the conclusion that the mechanosensation-dependent departure of PC1 from the primary cilium is due to ATP secretion and consequent PKA activation, which lead to PLAT domain phosphorylation of PC1 and its arrestin-dependent departure from the cilium.

**Figure 6.**
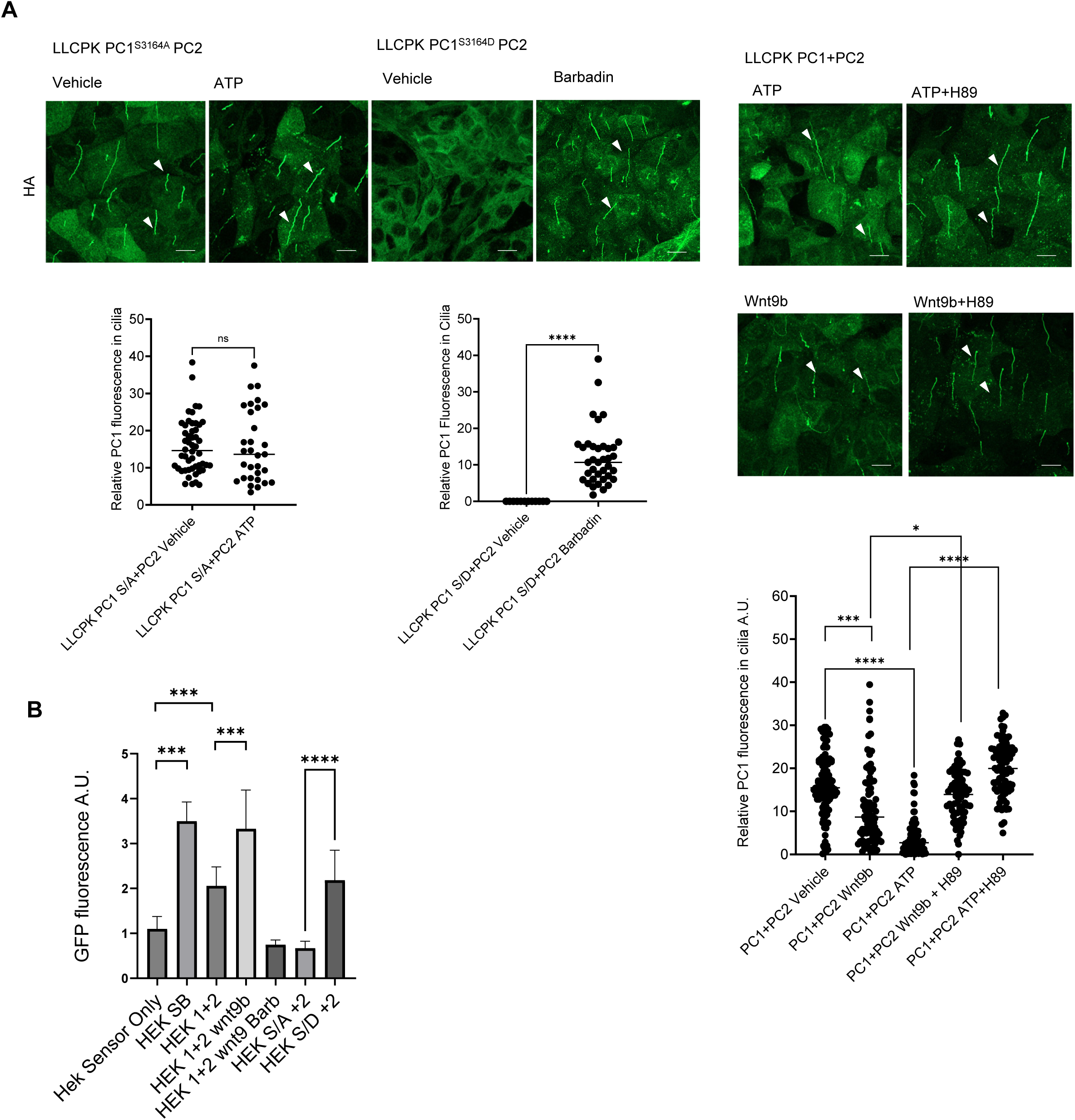
PKA is a key mediator of the signal-dependent modulation of the ciliary localization PC1. **(a)** Immunofluorescent detection of the localizations of PC1S3164A and PC1S3164D to the primary cilia in LLCPK cells. ATP treatment did not alter the ciliary distribution of PC1S3164A, whereas the ciliary exclusion of PC1 S3163D was reversed by Barbadin treatment. PKA inhibitor H89A treatment prevented the departure of PC1 from primary cilia that is induced by either ATP or Wnt9b treatment. **(b)** Biosensor assay was used to assess the importance of PC1 PKA phosphorylation site S3164 for PC1 activation-dependent inhibition of GSK3β kinase in HEK293 cells. Data are shown as mean ±SEM, n≥3 for all experiments. To assess statistical significance unpaired Student t-Test analysis was performed. P values <0.05 were considered significant *=P<0.05; **=P<0.01; ***=P<0.001.; ****=P<0.0001 Differences between mean values obtained for experimental conditions versus control conditions that produce minimal and/or maximal signals were evaluated. In panel 3d the values obtained for each condition were compared to those obtained in the GFP alone condition (indicated by the symbols above each bar) and pairwise comparisons of the values obtained in a subset of conditions related by a single experimental manipulation are indicated by the brackets above the bars.

To determine whether or how the effects of β-arrestin and PLAT domain phosphorylation influence the signaling capacity of PC1, we carried out the GSK3β fluorescent biosensor assay to measure PC1-induced GSK3β inhibition in HEK cells, as in Fig. 2c. We find that Wnt9b-induced, PC1 and 2-dependent inhibition of GSK3β is blocked by barbadin (Fig. 6b). In addition, the phosphomimetic S3164D PC1 protein, which is expected to interact constitutively with β-arrestin, induced substantial GSK3β inhibition in the absence of Wnt9b. These data strongly suggest that PLAT domain-mediated β-arrestin interaction^72^ plays a critical role in both the activation-dependent removal of PC1 from the cilium and the PC1 activation-dependent inhibition of GSK3β. These data are consistent with the possibility that, as has been shown for a number of classical GPCRs, β-arrestin not only mediates the internalization of activated PC1, but also plays an obligate role in downstream signaling mediated by activated PC1^56,73,74^.

If flow-induced mechanostimulation of primary cilia induces activation of PC1-dependent signaling pathways and the β-arrestin-dependent departure of PC1 from primary cilia, then we would expect that the shear force induced by fluid flow in the renal nephron would similarly result in exit of PC1 from cilia. To examine this possibility, we used the BAC transgenic mouse model (Pkd1^F/H^-BAC) that was employed in Fig. 2b, which uses the PC1 native gene structure to control the expression of a PC1 protein tagged with an N terminal 3XFLAG epitope and a C terminal 3XHA sequence ^35^. Immunofluorescence analysis employing anti-HA antibody to detect PC1, and anti-ARL13B to detect cilia, revealed little or no detectable PC1 in renal tubule epithelial cell cilia under basal conditions (Fig. 7a). We next performed a unilateral ureteral ligation to block renal tubule fluid flow for 30 minutes in one kidney of these Pkd1^F/H^-BAC mice. Immunofluorescence analysis revealed that this treatment resulted in the appearance of a readily detectable PC1 signal in cilia of renal tubule epithelial cells (Fig. 7a). In kidneys in which the UUO ligation had been released after 30 minutes, permitting tubule flow for an additional 30 minutes, no ciliary PC1 signal was observed (Fig. 7b). These data demonstrate that the mechanostimulatory effects on PC1 ciliary localization that we observed in LLC-PK1 cells are also detected *in situ* in nephrons in intact kidneys that experience normal renal tubule fluid flow. This suggests the surprising conclusion that, under normal circumstances, renal tubule fluid flow induces the departure of PC1 from the cilia. Localization of PC1 to cilia *in vivo* appears to be most detectable under conditions that interrupt tubule fluid flow.

**Figure 7.**
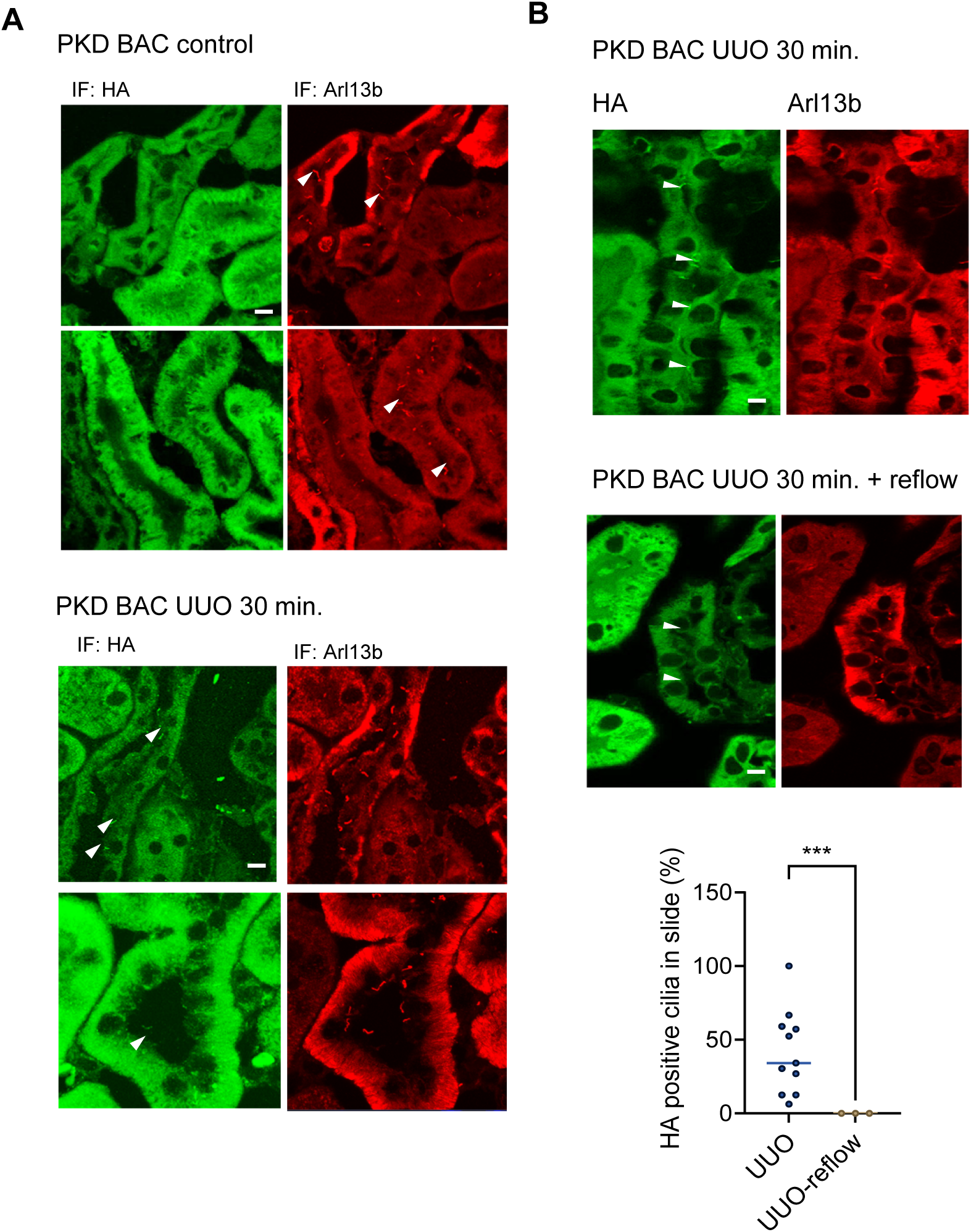
**Urine flow in mouse kidney renal tubules promotes removal of PC1 from the primary cilia of tubule epithelial cells**. **(a)** Immunofluorescent detection of the localization of PC1 (green) in PKD1-Bac transgene mouse kidneys that underwent the sham or temporary unilateral ureter obstruction for 30 min. Cilia are detected with an antibody directed against Arl13b (red) **(b**) Immunofluorescent detection of the localization of PC1 in PKD1-Bac transgene mouse kidneys subjected to UUO or UUO followed by release of the clamp. Ciliary detection of PC1 is detected only in those tubules in which flow is interrupted. Tubules in which flow has been interrupted and then reestablished lack PC1 staining in cilia. (Data are shown as mean ±SEM, n≥3 for all experiments. To assess statistical significance unpaired Student t-Test analysis was performed. P values <0.05 were considered significant *=P<0.05; **=P<0.01; ***=P<0.001.; ****=P<0.0001 Differences between mean values obtained for experimental conditions versus control conditions that produce minimal and/or maximal signals were evaluated. In panel 3d the values obtained for each condition were compared to those obtained in the GFP alone condition (indicated by the symbols above each bar) and pairwise comparisons of the values obtained in a subset of conditions related by a single experimental manipulation are indicated by the brackets above the bars.

## Discussion

The primary physiological roles of the proteins encoded by the genes mutated in ADPKD remain mysterious. The data presented here demonstrate that PC1 can function as a metabotropic receptor, and they uncover its role in the regulation of downstream effectors whose activities have already been linked to renal cystic disease. We find that Wnt9b activates PC1 through a mechanism similar to that employed by aGPCRs^19,75,76^. Like aGPCRs, PC1 undergoes autocatalytic cleavage within its GAIN domain to produce a large extracellular NTF that is non-covalently attached to the CTF that embodies the protein’s 11 TM domains^17,28^. Incubation with Wnt9b causes the PC1 NTF to dissociate from the CTF, exposing the “*Stachel*” at the N terminus of the CTF that acts as a TA to induce the protein’s signaling function. Removal of the NTF, either via Wnt9b binding or by exposure to alkaline pH, initiates assembly with and inhibition of the GSK3β kinase. As is the case with several aGPCR-mediated signals^43,46,51,53,77^, this process requires G_α13_ and leads to activation of RhoA, which in turn activates ROCK leading to phosphorylation of GSK3β at Ser9 and inactivation of its kinase activity. A version of PC1 that lacks the NTF behaves as a constitutively active receptor that maximally inhibits GSK3β independently of Wnt9b. In contrast, an identical protein that lacks the Stachel TA exerts no inhibitory effect on GSK3β, although this inhibitory activity can be partially recovered by adding back the Stachel TA as a soluble peptide. Similar observations have been made in an assay system that interrogates the activity of a different potential PC1 downstream signaling pathway^32^. The activity of GSK3β is elevated, and the levels of pGSK3β are reduced, in the pre-cystic and cystic kidney tissue from mice that manifest inducible, kidney-specific disruption of PC1 expression^10,11^. Furthermore, we find that the localization of PC1 to the primary cilium appears to be modulated by the activation of the PC1 receptor-like function. Exposure to Wnt9b causes PC1 to depart the cilium, and this departure appears to be dependent upon the involvement of β-arrestin. Similarly, in the absence of its NTF, PC1 is excluded from the cilium through a mechanism that requires the involvement of both its TA and β-arrestin. Mechanical stimuli cause PC1 to depart the primary cilium both *in vitro* and in cultured cells. This effect appears to be driven by mechanical stimulation-induced ATP secretion and is mediated by β-arrestin and PKA. Finally, the association with β-arrestin appears to play a critical role in the aGPCR-like effect of PC1 on the suppression of GSK3β activity. These data together define a new functional role for PC1 that places it in the center of a signaling network that is implicated in the pathogenesis of renal cystic disease.

Several lines of evidence fuel the long-held suspicion that PC1 can function as a receptor. The PC1 cytoplasmic C terminal tail interacts with several trimeric G proteins^12–15^ and a naturally occurring disease-causing mutation perturbs this interaction, suggesting that it is critically required for PC1 function^12^. PC1 protein undergoes an autocatalytic cleavage that is prerequisite for the separation of its NTF from the membrane-associated CTF^17^ at the GPS of its GAIN domain, which is highly homologous to those found in aGPCRs^18^, suggesting that PC1 may be an “atypical aGPCR”^28–32^. PC1 NTF binds to Wnt9b and Wnt3a with very high affinity (K_D_ in the range of ∼4 nM)^33^. In addition, recent studies indicate that the released NTF, or fragments of it, can act as a soluble ligand that activates the channel activity of the PC1, PC2 complex^78^. Our studies show that Wnt9b induces dissociation of the NTF and activates a G protein-dependent signaling cascade. These observations confirm that PC1 can participate in a functional aGPCR-like mechanism, transducing an interaction with a ligand to modulate a downstream effector that has been implicated in the pathogenesis of ADPKD^11^.

Because the recently solved structure of PC1 together with PC2^4^ was obtained using a PC1 construct lacking the NTF and the cytosolic C terminal tail, it does not provide spatial insights into the interactions and mechanisms that we describe. It is clear, however, that the PC1 protein does not resemble the canonical structure of a GPCR, either in the number and organization of its transmembrane segments or in the conformations of its cytosolic loops. The G protein-dependent signaling pathway that we have elucidated must, therefore, be considered in the context of the non-canonical structure of its receptor. Since the cytoplasmic C terminal tail of PC1 interacts directly with G proteins, and since this interaction appears to be a critical determinant of PC1 function *in vivo*, it is possible that the aGPCR-like process suggested by our data is mediated entirely through PC1^12–15^. According to this model, Wnt9b-induced dissociation of the NTF from the CTF reveals the tethered agonist, which engages with a binding pocket at the extracellular face of PC1’s 11TM domain, inducing a conformational change within the PC1 cytosolic tail that stimulates the binding and activation of G_α13_ via a mechanism analogous to but perhaps structurally distinct from that of traditional GPCRs. Recent modeling studies have suggested mechanisms through which the putative PC1 tethered agonist can functionally interact with residues on the extracellular-facing loops of the PC1 protein^31,32^. It is also possible, however, that Wnt9b binding induces assembly of a multi-molecular PC1-associated signaling complex that may incorporate one or more additional GPCRs, whose activity may then be modulated by the inclusion of PC1 in this complex. Interestingly, the aGPCR ADGRA2/GPR124 participates in a complex with other components of the Wnt pathway including Frizzled-type GPCRs to regulate canonical Wnt signaling induced by Wnt7^79–83^. Similarly, co-assembly with a purinergic receptor could contribute to the observed effect of extracellular ATP on the ciliary localization of PC1. Consistent with this idea of activation-induced assembly of a PC1-supercomplex is our observation that Wnt9b binding induces rapid formation of a complex that includes PC1 and GSK3β. Furthermore, a previous analysis of the G protein binding specificity of PC1 indicates that it can interact with high affinity with G_αs_, G_α14_, G_α12_, G_αi1_ and G_αi2_, but not with G_α13_^15^. Thus, our data demonstrating Wnt9b-enhanced co-precipitation of PC1 and G_α13_ may suggest that activation of the receptor function of PC1 induces its assembly with another protein that interacts with G_α13_, a G protein that is activated by a number of aGPCRs^50^. Our experiments employing PC1^ΔNTF^ and PC1^ΔNTF+ΔTA^ demonstrate that whatever type of signaling complex assembly may occur requires the presence of the tethered agonist. Finally, it has been suggested that PC1 may act as a negative regulator of G protein activity, perhaps by serving as a scavenger of G protein subunits that prevents them from participating in signaling processes^15,84^. Whether and how either of these possible mechanisms relates to PC1-mediated G protein activation will be determined in future experiments designed to define the components of the Wnt9b-induced PC1-containing complex in full and to identify the structural determinants within the PC1 protein itself that are required for this transduction to occur. When the full length PC1 protein is employed in the fluorescence-based GSK3β activity assay ^37^, Wnt9b-induced kinase inhibition only occurs if the PC2 protein is co-expressed. Since cell surface delivery of PC1 appears to require co-expression with PC2^47^, our finding is consistent with the natural supposition that PC1 must be present at the plasma membrane to interact with extracellular Wnt9b. Interestingly, however, the PC1^ΔNTF^ construct does not exhibit any requirement for PC2 co-expression in order to inhibit GSK3β or to be delivered to the cell surface. These observations suggest that, in at least a subset of PC1-dependent signaling processes, PC2 is required for the proper maturation and trafficking of PC1 rather than as an obligatory component of the receptor machinery. Co-expression with PC2 facilitates the autocatalytic proteolytic cleavage of PC1 at its GAIN domain, and this cleavage is required for the forward trafficking of the PC1 protein to the cell surface^47–49,85^. Our data suggest that, in the absence of the NTF, the PC1 CTF can traffic and function independently of any need to associate with PC2. Since the localizations of PC1 and PC2 are overlapping but not identical^3^, our data support the suggestion that PC1 that does not co-localize with PC2 may nonetheless be able to mediate signal transduction. Interestingly, a previous study reported that PC1 can activate G proteins and that this activity is suppressed by co-expression with PC2, consistent with the possibility that G protein signaling mediated by PC1 involves a population of this protein that is not permanently associated with PC2^86^. Finally, some or all of the Wnt ligand-mediated activation of PC1 might occur in intracellular compartments rather than at the cell surface. According to one iteration of this model, newly-synthesized Wnt ligand could act as an autocrine activator of signaling mediated by newly-synthesized PC1 while they traffic together through the secretory pathway. Examples of such intracellular signaling phenomena have been observed for other GPCRs^87^. It is also worth noting in this context that we find that the capacity to interact with β-arrestin appears to play a critical role in the suppression of GSK3β activity that is induced by PC1 activation, as is the case for a number of GPCR-mediated signaling processes that occur in intracellular compartments^56,73,74^.

Both PC1 and PC2 localize at least in part to the primary cilium^7,88^ and their presence there appears to be critically important for some aspects of their signaling function. Cilia that lack functional polycystin proteins produce an as yet uncharacterized cilium-dependent cystogenic activity that is suppressed by the presence of the polycystin proteins^38^. The data presented here demonstrate that ligand-induced activation of the receptor functions of PC1, as well as ciliary mechanosensation, induce the β-arrestin dependent departure of PC1 from the cilium. Consistent with previous studies^72^, we find that this β-arrestin-mediated activity-dependent departure of PC1 from primary cilia requires the activity of PKA and is critically dependent upon the phosphorylation state of residue S3164 in the PLAT domain of PC1. It remains to be determined whether the receptor-like activities that we describe, and their capacity to inhibit of GSK3β kinase activity and modulate PC1 ciliary localization, are mechanistically related to the PC1-dependent suppression of the cilium-dependent cystogenic activity. It is interesting to note in this regard that GSK3β has been shown, at least in certain cell types and circumstances, to localize to cilia or basal bodies ^89,90^ and to play a role in regulating cilia formation and stability^91^. It is also worth noting that cilia sense nutrient levels and communicate with mitochondria to regulate metabolism^92^,and that polycystins play direct roles in modulating mitochondrial function^8,93–96^. It remains to be determined whether the processes that we have identified may link ciliary polycystin activity to the cilia-mediated effects on mitochondrial function.

Our data suggest that, under conditions of normal renal tubule fluid flow *in vivo*, the PC1 protein may be fully activated, as evidenced by its relative absence from the primary cilium of renal tubule epithelial cells and by the increased accumulation of PC1 in cilia when renal tubule fluid flow is prevented. Taken together, these observations are consistent with the surprising possibility that PC1-dependent signaling, and its presumed capacity to suppress CDCA, are associated with processes that drive PC1 out of the cilium rather than with processes that enhance its ciliary accumulation. It is worth noting in this context that a large number of papers report that alterations in ATP secretion, in the ATP secretory machinery, and in purinergic receptor expression and activation, are detected in ADPKD cyst epithelial cells and may contribute to cyst development or expansion ^97–106^. Furthermore, shear-induced bending of the primary cilium stimulates ATP release from renal epithelial cells *in vitro* and *in vivo*^69–71^. Thus, it is possible that the ATP-induced relocalization of PC1 out of the cilium that we have observed may reflect a putative role for PC1 in responding to the presence of extracellular ATP and in initiating processes that suppress the pro-cystogenic activity of this ATP. This model posits that ATP secreted in response to ciliary detection of shear stress may account for or help to drive CDCA, that PC1 is directly or indirectly activated by this ATP signal and, as a result of this activation, that PC1 contributes to suppressing the cystogenic influence of flow-induced ATP release. Future experiments will be required to assess the validity of this hypothesis and of the possible relationship between CDCA and the receptor-like properties of PC1 that we have observed. Finally, it should be noted that while we have shown that Wnt9b, shear stress and ATP signals can activate the receptor function of PC1 and that G_α13_, RhoA, ROCK, GSK3β, PKA and β-arrestin can participate in the PC1-dependent response to these signals, it remains quite possible that other ligands or mechanical processes may stimulate the PC1 receptor and that other effectors may participate in transducing its signals ^31,32^. The tools that we have developed will facilitate future efforts to interrogate the upstream and downstream arms of the PC1 receptor signaling pathway and to explore novel therapeutic strategies based upon manipulating its activity.

## Methods

### Antibodies

Antibodies used: anti-HA (Roche, 118 674 23 001); anti Arl13b (UC Davis/NIH Neuro mAB 75287) anti-FLAG, anti-Actin (F7425, A2228, Sigma); anti-GSK3β, anti-RhoA (12456T, 2117S Cell Signaling); anti-pS9GSK3βand Gα13 (ab107166, ab128900 Abcam); anti C-terminal PC1 (EJH002 Kerafast); For Li-Cor Western Blot scanning, IR-680 and –800 dye-conjugated antibodies from Li-Cor (926-6708, 926-32211) and streptavidin IR-dye conjugates (Li-Cor) were used as secondary reagents. For microscopy, Alexa dye-coupled antibodies (Alexa-488, 594, 647; Molecular Probes) were used as secondary reagents.

### Plasmids

The PCDNA3.1 zeo vector constructs encoding PC1 and PC2 were described previously^34^. The PC1^ΔNTF^ construct was produced synthetically by Twist Biosciences and provided in the pTwist vector (Twist Biosciences, South San Francisco, CA) (full sequence available on request). In brief, the PC1 CTF sequence, beginning with the TAF residues at its N terminus, was preceded by sequence (MYPYDVPDYAQAVDNLTSAPGNTSLCTRDYKITQ) that includes an HA epitope tag followed by a portion of the human P2Y12 receptor to ensure membrane targeting of the PC1 construct^28^. The PC1^ΔNTF+ΔTA^ construct was generated by excising the Stachel sequence (TAFGASLFVPPSHVQFIFPEPSAS) from the PC1^ΔNTF^ construct using flanking restriction sites (sequence available on request). PC1^ΔGBS^ has been described previously^34^. In the nomenclature employed in this previous publication, PC1^ΔGBS^ was referred to as PC1^ΔNLS^. The chimeric GFP-GSK3β phosphorylation site construct was prepared according the previously published description ^37^ PC1^S3164A^ and PC1^S3164D^constructs were generated as previously reported^72^ (Prof. Albert Ong, University of Sheffield, UK).

### Peptides

Oligopeptide fluorescent-dye conjugates for kinase gel shift assays were synthesized by GenScript (Piscataway, NJ) according to the previously described design^39^. Peptide A (TAFGASLFVPPSHVQFIFPEPSAS) corresponding to the full-length PC1 Stachel sequence and scrambled Peptide A (FLFEFSPAFIPSGSPSQATVHVPA) were synthesized by GenScript.

### Cell lines and transfection

HEK293 cells (human female origin) were cultured as described ^47^ in DMEM supplemented with 10% fetal bovine serum, 1% penicillin/streptomycin, and 1% l-glutamine. The Pkd1^+/−^ Het cell line and the Pkd1^−/−^ Null cell lines correspond to the PH1 and PN24 lines, respectively according to the nomenclature employed in their original description^41^. These cells were maintained and passaged in DMEM/F12 supplemented with 10% FBS and γ-interferon (5 U/ml; Sigma-Aldrich, St. Louis, MO, USA) at 33 °C and 5% CO2. To inactivate the immortalizing transgene and thus initiate differentiation, cells were transferred to γ-interferon-free medium and maintained at 37 °C for 5 days before prior to their use in experiments^41^. The mouse renal tubule M113 cell line, which lacks PC1 expression, was maintained as previously described^36^.

Transient transfections were performed using HEK293 cells and Lipofectamine 2000 (Invitrogen). The quantities of the cDNA constructs used for transfection were adjusted to yield similar levels of protein expression as assayed by Western blot. Transfected cells were maintained for 24 to 48 hours post transfection before experimental analysis. Stably transfected cell lines were created by clonal selection of positively transfected cells and their propagation in MEM supplemented with antibiotic for selection pressure (Zeocin at 0.3mg/ml for PC2 expressing cells and G418 at 0.8mg/ml for cells expressing all of the versions of the PC1 constructs). LLCPK PC1+PC2 and LLCPK PC1^ΔGBS^+PC2 cells were generated as described in Gilder et al., 2018^34^. To ensure the development of a well established primary cilia, LLCPK cells were allowed to grow to confluency and then maintained for at least 4 days.

### Mechanical Stimulation Assay

To assess the effects of mechanical stimulation on polycystin protein localization to the primary cilium, LLCPK cells were plated in culture dishes, allowed to become confluent and fully polarized for at least 4 days, and then placed onto a horizontal shaker residing within a standard CO2 cell culture incubator. Depending on the size of the culture dishes, cells were exposed to different spinning speeds e.g. 24-well plates were spun at 100 rpm, 6-well plates at 38 rpm and 10cm dishes were spun at 25 rpm for desired periods of time.

### Mouse Line

The Pkd1^fl/fl^, Pax8^rtTA^, TetO-Cre mouse line ^38^ was employed to examine in the activity of downstream PC1 signaling effectors in mouse kidney cystic tissue. All experiments were performed according to protocols that were reviewed and approved by Yale University’s Institutional Animal Care and Use Committee. Each condition employed 3 female and 3 male mice. To induce kidney-specific Cre expression, 2 mg/ml doxycycline was added to the animals’ drinking water at postnatal day 28 and removed after 14 days. Mice were sacrificed either prior to doxycycline induction or at the 6 week or 12-week time points after the completion of the doxycycline induction. The transgenic animals that express Flag-PC1 3xHA-Bac construct were the kind gift of Dr. S. Somlo (Yale University) and their production has been previously described^35,48^

### Short term unilateral/bilateral ureter obstruction (UUO/BUO) to study effect of renal apical flow on PC1 ciliary localization

For the short term UUO/BUO procedure, animals were anesthetized with ketamine/xylazine mixture at 10mg/kg. The abdominal cavity was exposed and ureter isolation performed using fine surgical forceps (Fine Science Tools cat # <u>00108-11</u>). Temporary clamp of the right and/or left ureter was performed using vascular clamps (Fine Science Tools cat # <u>00400-03</u>). During the 30 min. clamp the surgical field was covered with gauze soaked in PBS to prevent the local dehydration of the tissue. After the ureter clamp, animals were sacrificed by intracardiac total body prefusion with ice-cold PBS followed by ice-cold fixative (4% PFA in PBS). Fixed kidneys were embedded in O.C.T. (Tissue-Tek cat# 4583) and processed for cryo-sectioning and microscopic analysis.

### Immunofluorescence analysis of cultured cells and of mouse renal tissue

HEK293 cells grown on poly-L coated coverslips were fixed (4% PFA in PBS) and permeabilized (PBS, 2% BSA, 0.1 % Triton X100). LLCPK cells were allowed to become fully confluent, after which they were maintained in culture for at least 96 hours prior to MetOH fixation^34^. Kidney tissue was fixed via whole body ventricular perfusion with PBS, pH 7.4 for 15 s followed with freshly prepared 4% formaldehyde in PBS for 2 min. Kidneys were removed, dissected into transverse slices 1 mm thick, and post-fixed for 1 h at room temperature, followed by three 15-min washes in PBS and cryomicrotome sectioning to produce 4 μm sections. Kidney sections were subjected to 15 min of antigen retrieval through incubation in 10 mM, pH 6.0 citrate buffer. Following multiple washes with PBS and quenching with 0.5 M NH4Cl, sections were washed with 1% SDS and blocked with 0.1% BSA in 10% goat serum buffer (blocking buffer).

For labeling of tissue sections and coverslip-grown cells, primary antibodies diluted in blocking buffer (1:300) were applied for 1 hour at RT, followed by several PBS washes and incubation with secondary antibody (1:200) diluted in blocking buffer. Following several additional PBS washes, coverslips were mounted on slides with VectaShield (Vector Laboratories, Burlingame, CA). Samples were imaged using a Zeiss LSM780 and LSM900 confocal microscopes according to previously published imaging parameters to ensure that image intensity could be quantitated ^34,47^.

### Cell Surface immunofluorescence

Cell surface immunofluorescence was performed according to previously published methods^34,47^. Briefly, cells were grown to confluence on poly-L coated coverslips, washed once with 4°C PBS and treated with anti-Flag antibody (1:300) in PBS with 1% BSA for 1 hr at 4°C. Cells were then fixed (4% PFA in PBS, 30 min, 4°C), washed twice with PBS and permeabilized with PBS containing 2% BSA, 0.1% Triton X100. Primary Rat anti-HA or Rabbit anti-Flag antibodies were applied (1:300) for 1 hr at RT or overnight at 4°C followed by three washes with PBS and 1hour incubation with secondary anti-rabbit Alexa 594 and anti-rat Alexa 488 (1:200). Following three washes in PBS the signal was visualized using a Zeiss LSM780 confocal microscope (63X objective, 1.4 NA) according to previously published imaging parameters that are designed to ensure that image intensity can be quantitated accurately^34,47^.

### Immunoprecipitation

Immunoprecipitations were performed from renal tissue lysates or lysates of HEK293 cells transiently or stably transfected to express PC1 and PC2 or various mutated forms of PC1. For total protein lysate preparation kidneys were harvested after PBS perfusion. Tissue samples were processed in brinkman homogenizer and resuspended in IP lysis buffer followed by protein extraction steps. Similarly, cell pellets were, washed with PBS, and resuspended in IP lysis buffer (50 mM Tris pH 7.4, 100 mM NaCl, 0.5% NP-40, 0.5% Triton X100, 1 mM Ethylenediaminetetraacetic acid (EDTA)). Both tissue and cell materials were sonicated for 60s, (single continuous 15s bursts at 40% power separated by 5s each) and incubated on ice for 1hr to ensure complete protein solubilization. Lysates were centrifuged (10,000xg, 10 min) to remove insoluble material. Anti-HA-conjugated dynabeads (Pierce/Thermo, 88837), equilibrated in lysis buffer (30 μL beads per reaction in 1 ml of the lysis buffer) for 10 min at RT on a rocking shaker, were incubated with cell lysates for at least 30 min at RT or overnight at 4°C. Following three washes with 1ml lysis buffer, dynabeads were magnetically recovered and precipitated proteins were eluted with SDS-PAGE loading buffer at 37-95°C for 10 minutes.

### Western blotting

Cell lysates or immunoprecipitated proteins were separated on a 10–12% SDS-polyacrylamide gel and electrophoretically transferred to nitrocellulose (Bio-Rad, 1620115). Quenching incubation with blocking buffer (PBS, 7% (w/v) powdered milk/BSA, 0.1% Tween) for 60 min at room temperature was followed by incubation with primary antibody at 4°C overnight: (monoclonal anti-HA (Rat) antibody (1:1000, Roche), polyclonal anti-FLAG (1:1000, Sigma), polyclonal anti-cMyc (1:2000, Sigma) or other antibodies as specified in the text). Subsequently, primary antibody binding was detected using species-specific infrared (IR)-conjugated secondary IgG (1:5,000−1:10,000; Li-Cor). Membranes were visualized with an Odyssey Infrared Imager (Li-Cor Biosciences) and quantitated using the integrated software package.

For western blotting experiments employing mouse renal tissue, crude kidney lysates were prepared by homogenization of snap frozen kidney tissue. Homogenization was performed using a Brinkmann mechanical homogenizer in IP Tris lysis buffer (50 mM Tris pH 7.4, 100 mM NaCl, 0.5% NP-40, 0.5% Triton X100, 1 mM EDTA) on ice for 15 seconds at 300 rpm. Homogenates were then sonicated for 60s, with single continuous 15s bursts at 40% power separated by 5 s each and left on ice for an hour to complete protein solubilization. Lysates were centrifuged (10,000xg, 10 min), resolved by SDS PAGE, transferred to nitrocellulose and analyzed by western blotting.

### Cell surface biotinylation

Surface labeling of the NTF or of the CTF of PC1 was performed according to standard cell surface biotinylation methods^107^. Following surface biotinylation, cells were lysed using standard cell lysis buffer (50 mM Tris pH 7.4, 100 mM NaCl, 0.5% NP-40, 0.5% Triton X100, 1 mM EDTA) with protease and phosphatase inhibitors (Roche, 4906845001, 11697498001). For assessment of cell surface-bound PC1 NTF, the biotinylated proteins were precipitated with streptavidin-conjugated agarose according to published methods^107^. Anti-Flag antibodies produced in Rabbit (Sigma, cat #F7425) were used (1:1000) in subsequent western blot analysis to detect the surface-exposed fraction of the N-terminal portion of PC1. To assess the biotinylation of the CTF, lysates of biotinylated cells were subjected to immunoprecipitation with Anti-HA-conjugated dynabeads (Pierce/Thermo, 88837) as described above. Recovered proteins were separated by SDS-PAGE and electrophorectically transferred to nitrocellulose, after which they were blotted with IRDye 800CW streptavidin conjugate. Signal was detected using Li-Cor Odyssey infrared imaging system.

### Analysis of cell signaling pathway activities

*Kinase activity gel shift assay:* Detection and quantification of GSK3β kinase activity in lysates of mouse renal tissue samples was performed using a non-radioactive *in vitro* kinase assay adapted from previously published methods^39^. For this *in vitro* kinase assay, recombinant active GSK3β kinase was obtained from Abcam (ab60863). Briefly, the *in vitro* kinase reaction was assembled by adding together recombinant purified GSK3β kinase and a custom made GSK3β target oligopeptide fluorescent dye conjugate (10μM in 30 μL reaction). Depending on the experimental conditions, reactions were additionally supplemented with crude lysates from un-induced or doxycycline-induced Pkd1^fl/fl^, Pax8^rtTA^, TetO-Cre mouse kidneys. The reaction was buffered using the standard NEB T4 ligase buffer and incubated at 30°C for 1 hr. To visualize the results of the kinase reaction, samples were separated on a 3% agarose gel followed by fluorescent imaging with GelDoc trans-luminometer.

*Quantitative GSK3β kinase fluorescent biosensor assay:* Measurement of endogenous GSK3β kinase activity was performed in HEK293 cells that were transfected with a construct encoding a chimeric GFP-GSK3β phosphorylation site fluorescent biosensor protein^37^. HEK293 cells were seeded in 24-well or 96-well poly-L lysine coated tissue culture plates and maintained in DMEM cell culture medium containing 10% fetal bovine serum, L-glutamine, and penicillin/streptomycin. HEK293 cells were transfected using Lipofectamine 2000 (Thermo Fisher, <u>11668019</u>). Transient co-transfections with plasmids to be tested (encoding PC1, PC2 or various forms of PC1^ΔNTF^) and the biosensor plasmid were performed in replicates of six. Recombinant Wnt9b (0.5ng/μL) (R&D Systems, 3669) was applied for 3 hrs or overnight. Each experiment included a set of wells that was treated with a chemical inhibitor of the GSK3β kinase: SB 266763, 10μM (Sigma, A2228), which served as a positive control. Plates were incubated at 37°C and 5% CO_2_ for the desired period of time. After incubation, the DMEM was replaced with PBS and GFP fluorescence was assessed using the Tecan plate reader (Infinite M1000).

*Activated RhoA assay:* Detection and quantification of the GTP-bound active form of RhoA in HEK cells was performed through precipitation using rhotekin-decorated agarose beads as previously described^55^. Briefly, a construct encoding the Rho binding domain of rhotekin fused to GST (Addgene, 15247) was employed for bacterial expression and purification. BL21 bacteria cultures were transformed with the RBD-GST plasmid, stimulated with 0.3 mM IPTG for 12 hrs at 16°C and lysed by sonication in a lysis buffer containing 1% Triton X100, 100mM NaCl and 25mM Tris and protease inhibitor cocktail, pH 7.5. Recombinant protein was recovered with glutathione agarose beads (Thermo Fisher, 16100) that were added to the lysate (50 μL of the beads slurry per 1 ml of the protein lysate). After 30 min incubation, beads were washed three times with the lysis buffer and used as bait in the RhoA-GTP pull-down assay. Briefly, RBD decorated beads were incubated with crude lysates from HEK1+2 and HEK PC1^ΔNTF^ cells prepared using the standard Tris lysis buffer described earlier. After three washes (with 1 ml of lysis buffer for each 50 μL of the beads), at 4°C, 10 min. for each wash, the beads were incubated with SDS-PAGE sample buffer and the recovered protein was resolved by SDS-PAGE followed by transfer to nitrocellulose and detection by western blotting using an antibody directed against RhoA (2117S Cell Signaling).

*RNAi mediated knockdown of the Gα13 and RhoA expression:* RNAi mediated knockdown of the G_α13_ and RhoA was performed using commercial RNAi cocktails (Santa Cruz, CA, anti-RhoA RNAi, anti-G_α13_ RNAi). For each experiment, cells were pre-transfected with 100nM of the corresponding RNAi oligos according to the manufacturer’s recommended protocol and allowed to recover for 24 hrs before the secondary transfections with cDNA constructs of interest were performed.

### Quantification and Statistical Analysis

Data quantification and plotting were performed using GraphPad software. Quantification of the results is presented as mean ± SEM. The N for each experiment is at least 3 and is noted in each figure legend. Differences between mean values obtained for experimental conditions versus control conditions that produce minimal and maximal signals were evaluated pairwise using the paired Student’s *t* test. Values of P<0.05 were considered to be significant. In addition, hypothesis-driven pairwise comparisons of the values obtained in a subset of conditions related by a single experimental manipulation were also performed.

### Summary of Supplemental Material

The Supplemental Material includes 5 figures that present additional data that provide support for the data depicted in each of the correspondingly numbered manuscript figures 1-5. Specifically, the Supplemental Material presents additional data relating to 1) PC1 GPS cleavage and surface biotinylation; 2) PC1 interaction with and inhibition of GSK3β; 3) Design, expression and activation of PC1 ΔNTF constructs; 4) Effects of PC1 activation on RhoA and G_α13_ activity and expression and distribution of PC1 ΔNTF constructs; and 5) effect of mechanical stimulation on PC1 ciliary localization

## Acknowledgments

We thank the Caplan laboratory for their valuable input and support. We also thank Dr. S. Thomsen (Vertex Pharmaceuticals) for helpful suggestions. We thank Dr. S. Somlo (Yale U.) and the Disease Models and Mechanisms Core of George M. O’Brien Kidney Center at Yale for providing the polycystic kidney disease mouse model and cell lines.

## Funding

This work was supported by NIH grants DK 120534, DK072612, DK 17433 (MJC), CDMRP PR093488 (W81XWH-15-1-0419) from the DoD (MJC), a Sponsored Research Agreement with Vertex Pharmaceuticals (MJC), and by grants from the Deutsche Forschungsgemeinschaft to N.S. (FOR2149 P01 [SCHO1791/1-2]; CRC1423 project 421152132, subproject B06) and T.L. (FOR2149/P01 [LA2861/4-2]; CRC1423 project 421152132, subprojects A06, B06).

## Author Contributions

N.G. designed, performed and interpreted the experiments and wrote and revised the manuscript; V.P., N.S. and T.L. contributed to experimental design and interpretation and reviewed the manuscript; M.J.C. designed and interpreted the experiments and wrote and reviewed the manuscript.

## Competing Interests

Michael J. Caplan has received research support and consulting fees from Vertex Pharmaceuticals.

## Data and materials availability

All data are available in the main text or the supplementary materials.

## Figure Legends

**Supp. Fig. 1.**
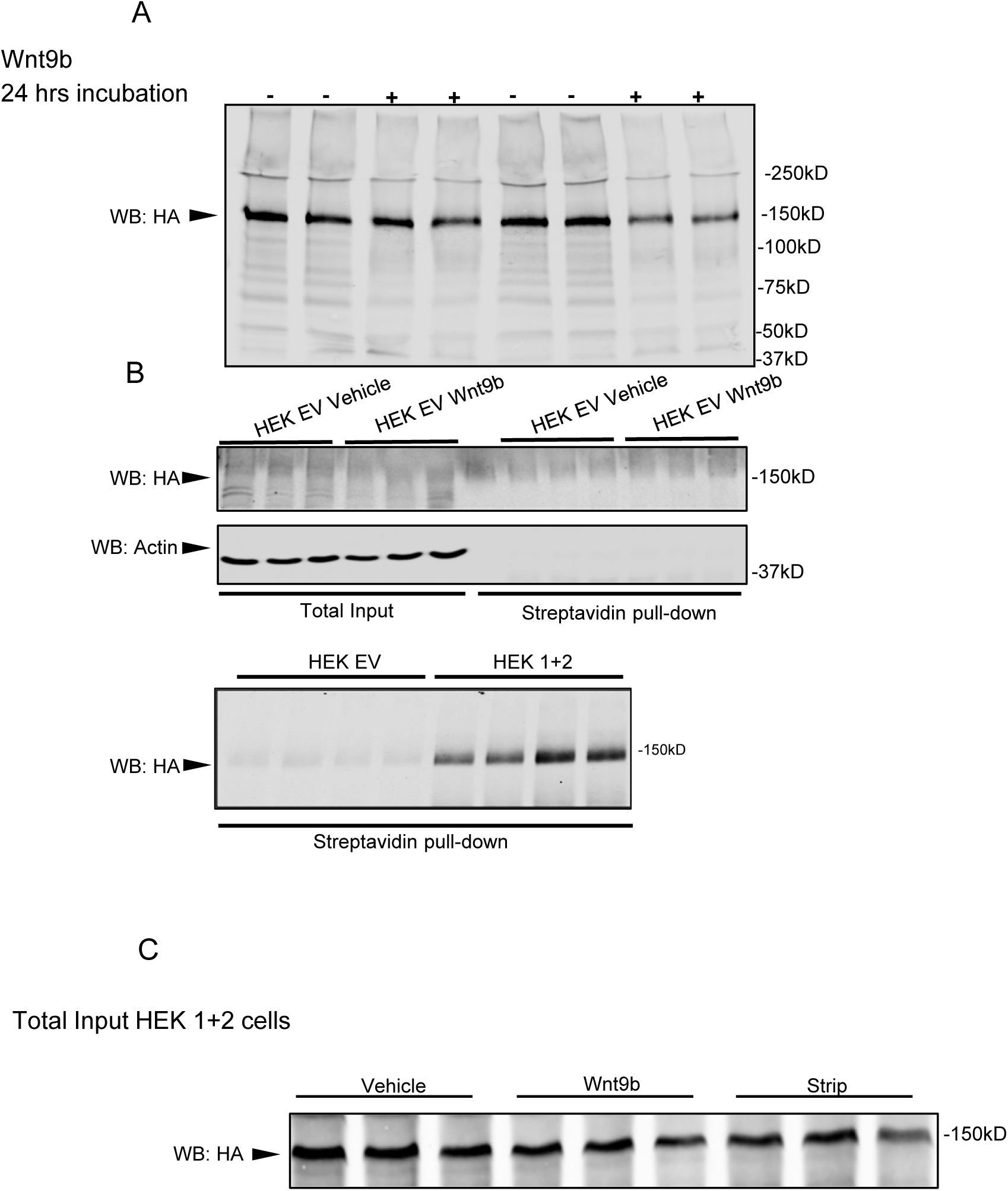
PC1 GPS cleavage, surface biotinylation and NTF shedding. **(a) GPS cleavage of the full length PC1 construct expressed in HEK293 cells is very efficient**. The panel depicts the anti-HA labeling of the entire western blot that is presented in Figure 1D. The PC1 protein employed in these experiments carries a Flag tag at its N terminus and a 3X HA tag at its C terminus. As evidenced by detection with both with anti-Flag in the main figure and with anti HA in this panel, the GPS cleavage of the expressed PC1 construct is extremely efficient. **(b) Controls for surface biotinylation experiments.** No protein in the 150 kDa weight range is detected by western blotting with anti-HA antibody in material recovered by streptavidin pull down from biotinylated HEK293 cells transfected with empty vector. Furthermore, a protein of 150 kDa is detected in material recovered by streptavidin pull down from HEK293 cells that express PC1. Thus, the surface biotinylation experiments presented in Figure 1 C and E specifically detect the PC1 protein**. (c) The quantity of cleaved CTF fragment of PC1 remained unchanged in cells that were used for the experiment depicted in** Fig 1 **E** Total HEK293 1+2 cell lysates were prepared after harvesting the MEM upon completion of WNT9b stimulation. Protein samples were analyzed by western blotting with anti-HA antibody N=3, statistical significance established by paired Student’s T test ***=P≤0.001; ****=P≤0.0001. It should be noted that the differing appearance of the NTF bands detected in Figure 1E versus HA bands in Supplemental Figure 1c (broad and fuzzy in Figure 1a, sharp in Supplemental Figure 1c) is in all likelihood attributable to different types of gel systems used to resolve these proteins in the two figures (gradient gel in Figure 1a, linear 10% acrylamide gel in Supplemental Figure 1c).

**Supp. Fig. 2.**
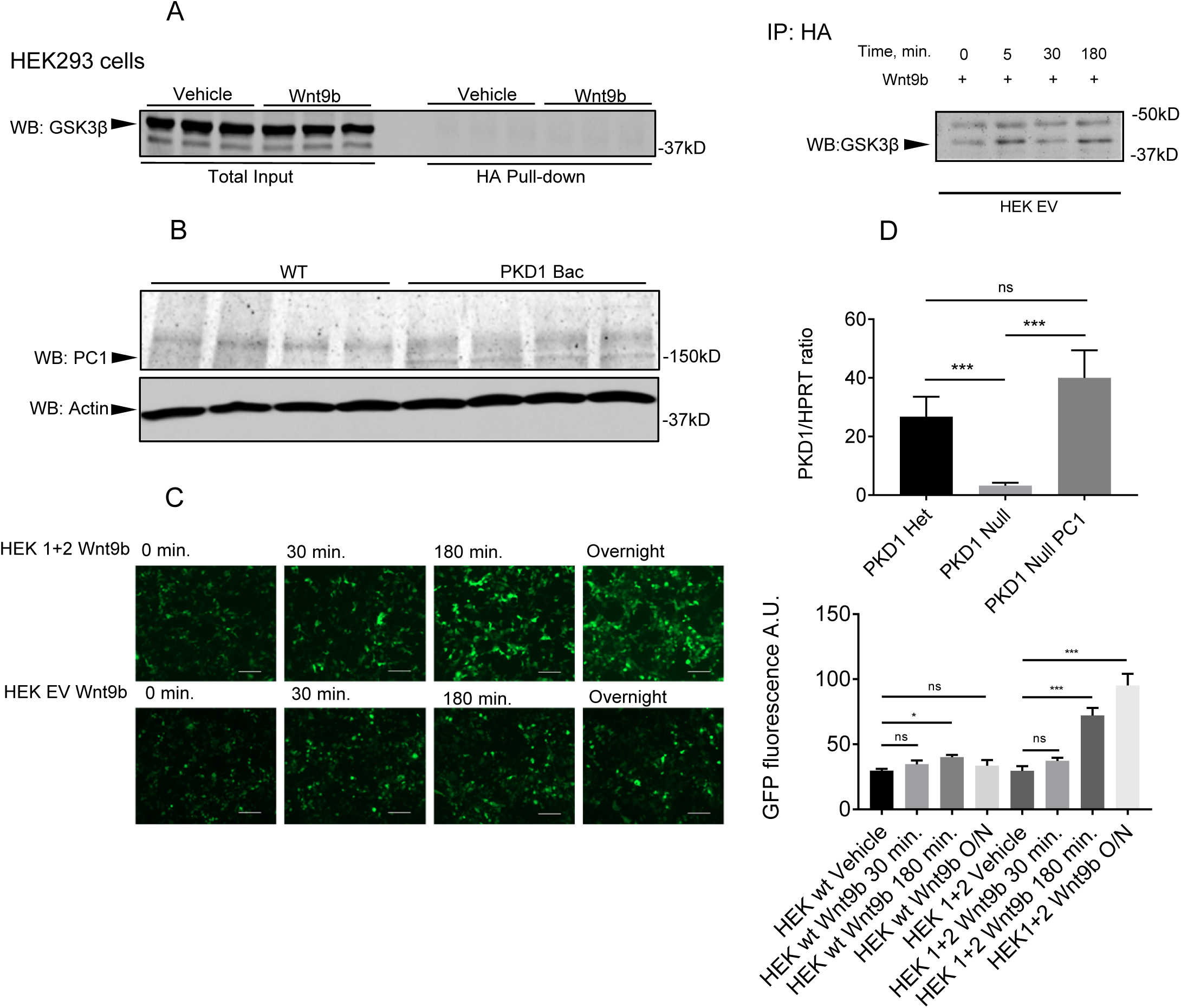
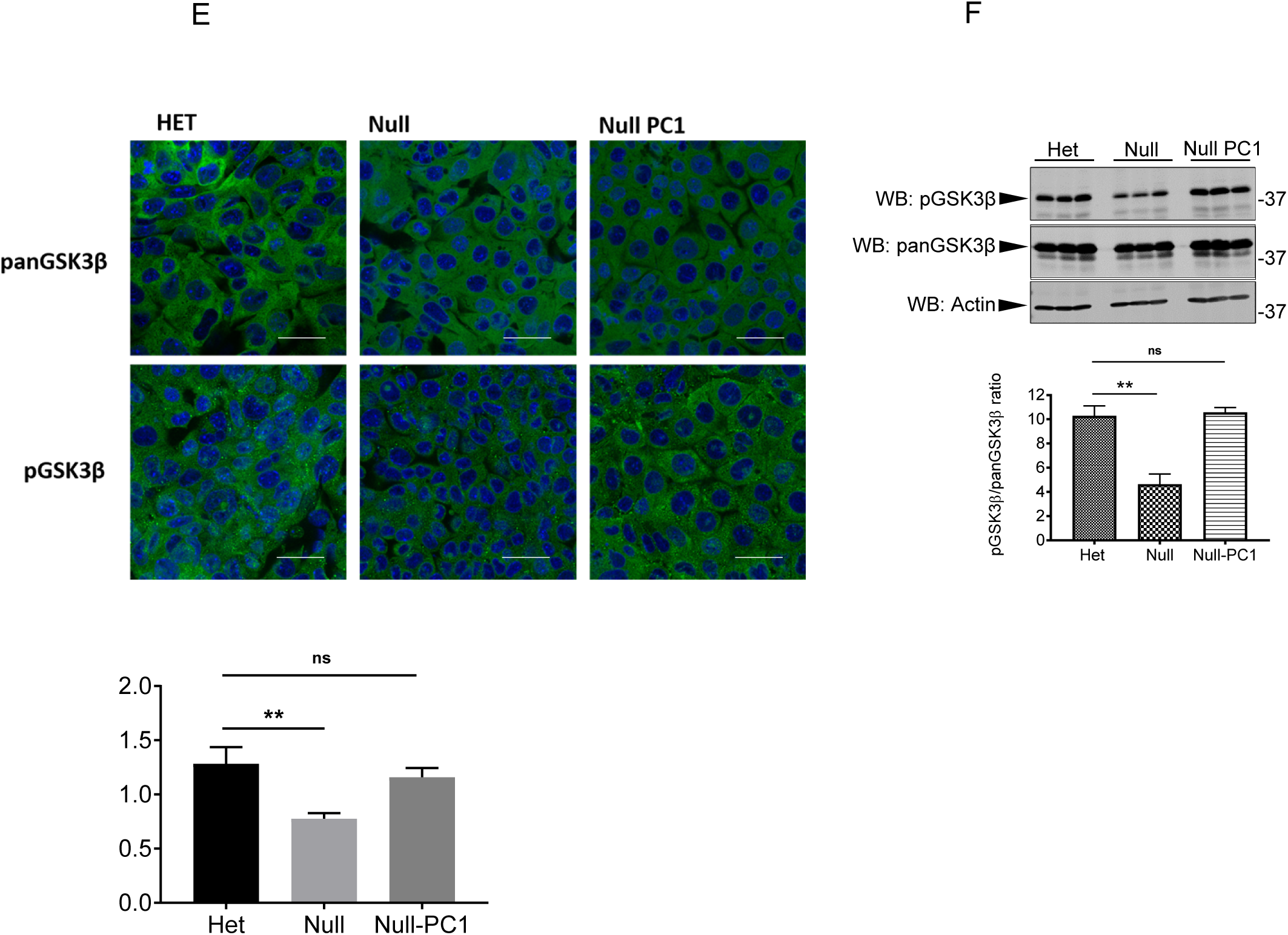
**(a) Wnt9b does not stimulate GSK3β precipitation with anti-HA antibody in cells that do not express PC1 and 2**. Lysates were prepared from HEK293 cells transfected with empty vector (EV) and treated overnight (left panel) or for the indicated number of minutes (right panel) with Wnt9b (13 nM). Lysates were subjected to anti-HA immunoprecipitation, separated by SDS-PAGE, electrophoretically transferred to nitrocellulose and probed with an antibody directed against GSK3β. Wnt9b treatment does not lead to an increase in the small quantity of GSK3β that is non-specifically precipitated by anti-HA antibodies in empty vector transfected cells that do not express PC1 and 2. n≥3 for all experiments. **(b) The PC1 CTF is recovered by immunoprecipitation with anti-HA antibody from lysates of Tg248 *Pkd1* BAC-transgenic mouse kidneys in which expression of PC1 carrying an N terminal FLAG tag and a C terminal HA tag is under the control of the native *Pkd1* promoter.** Lysates prepared from Tg248 *Pkd1* BAC-transgenic mouse kidneys were subjected to anti-HA immunoprecipitation and the recovered proteins were separated by SDS-PAGE and analysed by western blotting using an antibody directed against the PC1 C terminal tail. The 150 kDa band corresponding to the PC1 CTF is present in immunoprecipitates from the Tg248 *Pkd1* BAC-transgenic mice but not in lysates prepared from kidneys of wild type mice. **(c) Time course of Wnt9b-induced GSK3β inhibition in HEK293 cells expressing PC1 and 2.** Addition of Wnt9b to the medium bathing HEK293 cells expressing PC1 and 2 leads to a time-dependent inhibition of GSK3β activity, as revealed using the fluorescent biosensor assay of endogenous GSK3β kinase activity. HEK293 cells transfected with empty vector (EV) exhibit a substantially smaller response to Wnt9b. **(d) Detection of PKD1 messenger RNA in PKD1 Het PKD1 null and PKD1 null cells stably transfected with PKD1 construct.** mRNA isolated from immortalized mouse renal epithelial cells that are heterozygous for PC1 expression (Het), that are homozygous for the deletion of PC1 expression (Null), or that are homozygous for the deletion of PC1 expression and have been transfected to express PC1 (Null PC1) was analyzed by qPCR using primers designed to amplify mRNA encoding PC1 and the housekeeping gene HPRT. The ratio of PC1 to HPRT expression is depicted in the graph. N=3, statistical significance was assessed by paired Student’s T test. *= P<0.05, **= P<0.01, ***=P<0.001 **Quantification of GSK3β phosphorylation as a function of PKD1 expression. (e) Immunofluorescent detection of the endogenous pan– and phospho-GSK3beta in immortalized mouse kidney PKD1 HET PKD1 Null, and PKD1 Null-PC1 polarized cells.** Kidney cells were seeded onto 10 mm coverslips and allowed to grow to complete confluency for at least 4 days. Following the rounds of washing and staining with pan and phospho-specific antibody, the immunofluorescent analysis of the endogenous pan– and phospho-GSK3β was performed using laser scanning confocal microscopy. **(f) Western blot depicting the quantities of the pan– and phosoho-GSK3β protein expressed by PKD1 Het, PKD1 Null or PKD1 Null cells stably expressing PKD1 construct.** Cell lysates were prepared 24 hours after transfection, separated by SDS-PAGE and electrophoretically transferred to a nitrocellulose membrane. The blot was probed with an antibody directed against the HA epitope tag. N=3, statistical significance was assessed by unpaired Student’s T test. ***=P<0.001

**Supp. Fig. 3.**
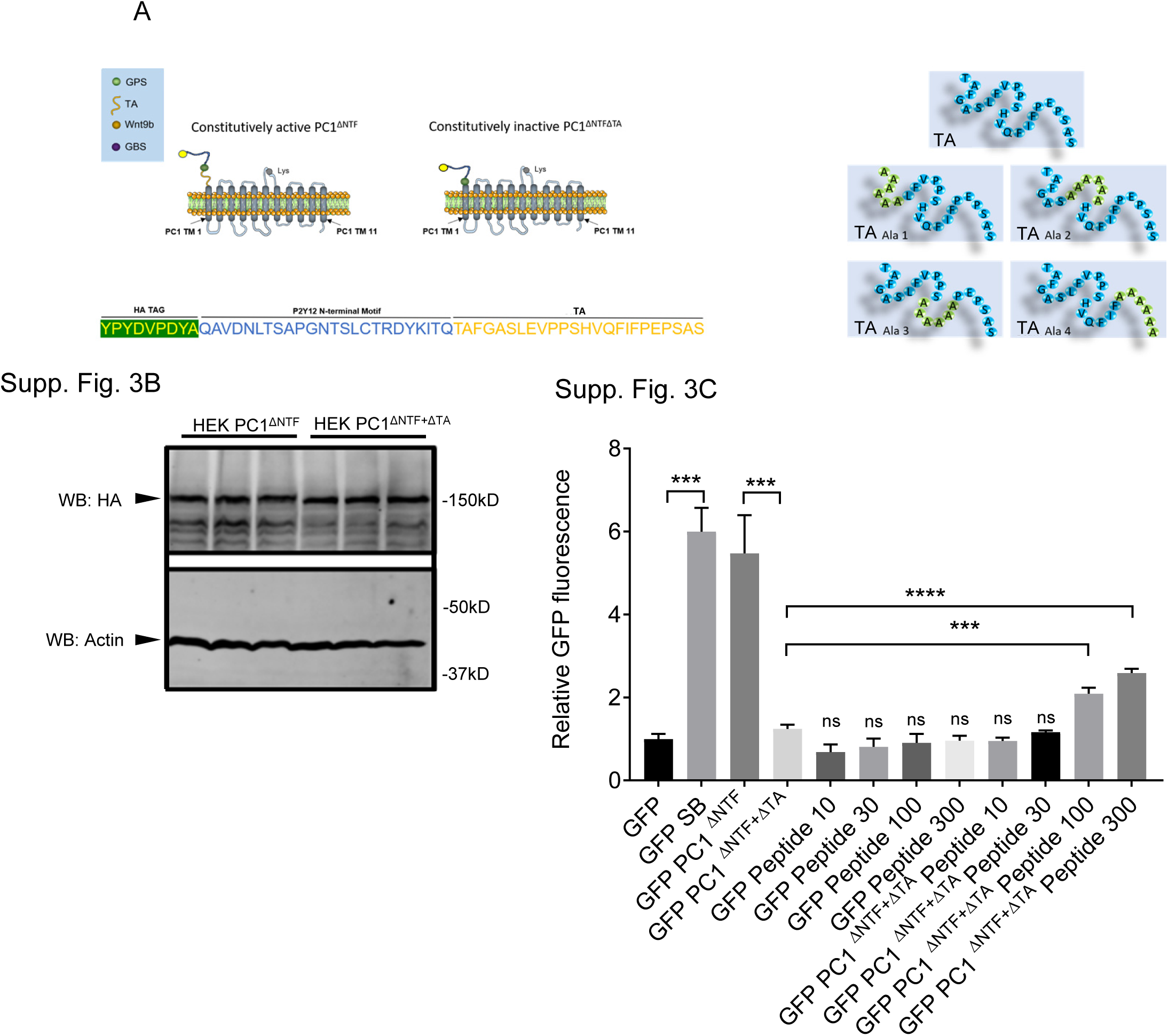
**(a) Schematic diagrams of PC1ΔNTF constructs**. Transmembrane domains are depicted in gray, the N terminal HA tag is shown in yellow, the P2Y12 sequence is drawn in black, and the tethered agonist (TA) (or stalk or “stachel”) segment appears in brown. The linear sequence of each of these pieces is shown in single letter code, and alanine substitution mutations have been inserted where indicated. **(b) Western blot depicting the quantities of the PC1^ΔNTF^ and PC1^ΔNTF+ΔTA^ proteins expressed by transient transfection of their encoding cDNA constructs in HEK293 cells.** Cell lysates were prepared 24 hours after transfection, separated by SDS-PAGE and electrophoretically transferred to a nitrocellulose membrane. The blot was probed with an antibody directed against the HA epitope tag, which is appended at the N terminus of both protein constructs, as well as with an antibody directed against actin, which served as a loading control. The PC1^ΔNTF^ and PC1^ΔNTF+ΔTA^ proteins are expressed at the same levels, supporting the conclusion that their differential effects on GSK3β kinase activity, as depicted in Figure 3B, is attributable to differences in their inherent signaling capacities. n≥3 for all experiments. **(c) concentration dependence of PC1^ΔNTF+ΔTA^ activation by soluble TA peptide.** The fluorescent biosensor assay of endogenous GSK3β kinase activity was performed on cells that express the GFP biosensor (GFP) plus PC1^ΔNTF+ΔTA^ or the GFP biosensor alone. Cells treated with the GSK3β inhibitor SB-216763 or transfected with the constitutively active PC1^ΔNTF^ construct serve as positive controls that define the maximal attainable signal. Cells expressing PC1^ΔNTF+ΔTA^ were incubated with 0, 10, 30, 100 and 300 μM concentrations of the soluble TA peptide. The ability of PC1^ΔNTF+ΔTA^ to inhibit GSK3β can be partially restored by addition of soluble TA (100 and 300 μM). This effect is not observed in cells that express the GFP biosensor alone. The graph depicts the means of the ratios of the GFP signals detected in each condition, normalized to the mean of this value obtained with cells transfected with the GFP biosensor alone. Data are shown as mean ±SEM, n≥3 for all experiments, Student t-Test or ANOVA multivariate analysis was performed where appropriate. P values ≤ 0.05 were considered significant.

**Supp. Fig. 4.**
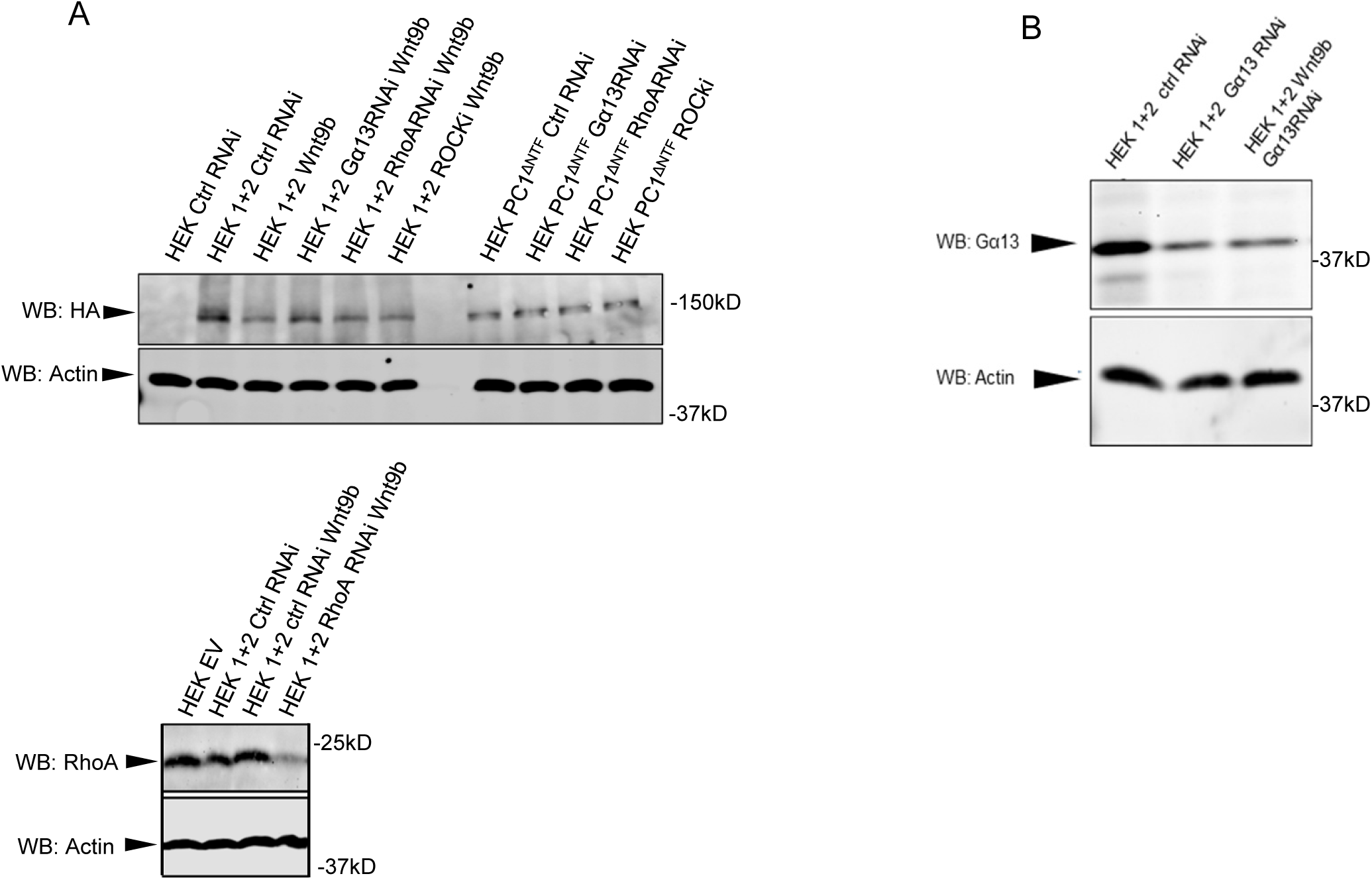
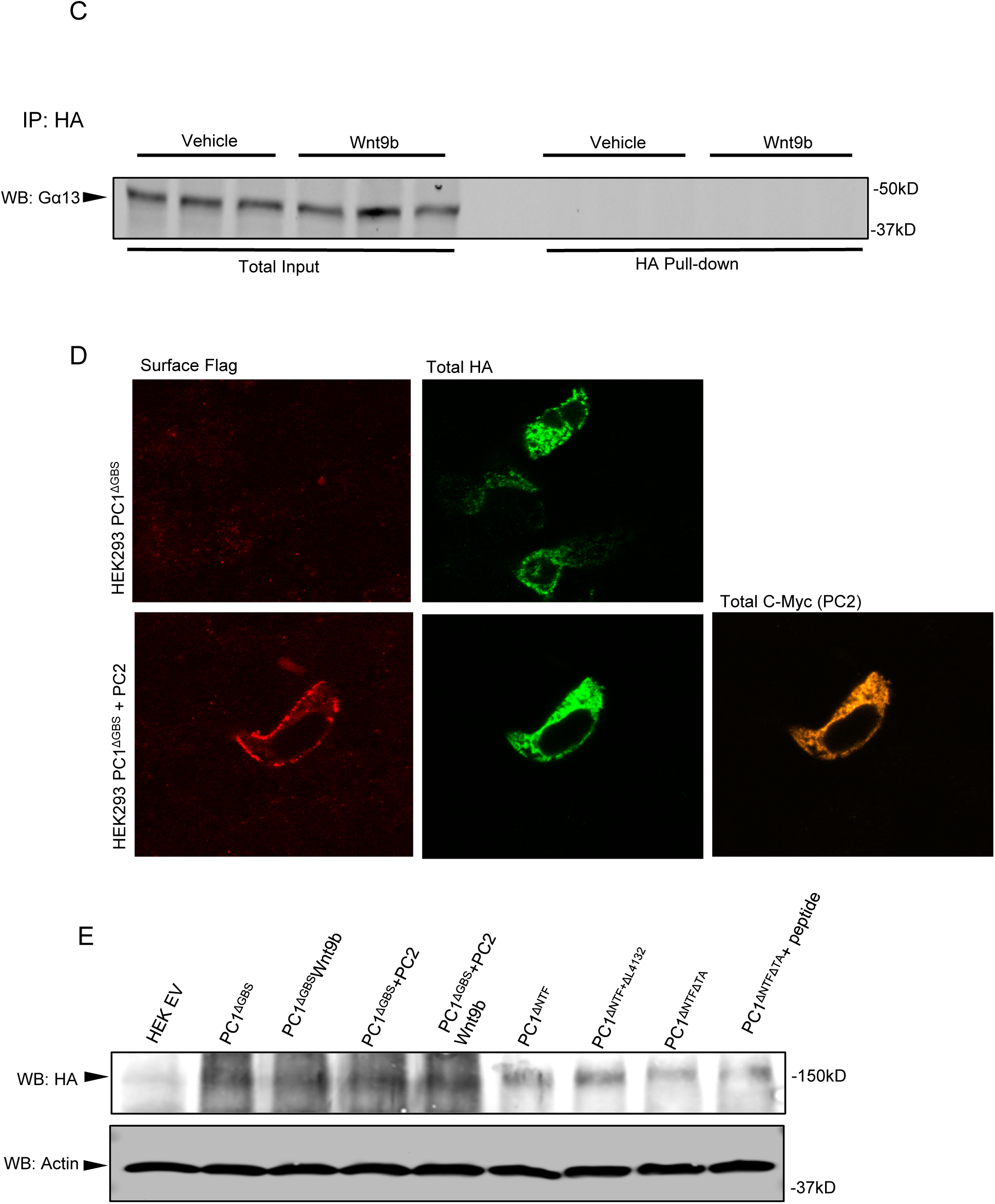
**(a) Assessment of expression of PC1 constructs and of RhoA in the samples employed in the fluorescent analysis of GSK3β activity depicted in Figure 4a**. *Upper left panel:* Lysates prepared from HEK293 cells subjected to each of the conditions analyzed in Figure 4a were separated by SDS-PAGE and analyzed by western blotting using antibodies directed against HA to detect the PC1 CTFand against actin as a loading control. *Lower left panel:* Cell lysates prepared 24 hours after treatment with RhoA-directed or control RNAi, were separated by SDS-PAGE and electrophoretically transferred to a nitrocellulose membrane. The blot was probed with an antibody directed against RhoA, as well as with an antibody directed against actin, which served as a loading control. Treatment with the RhoA-directed RNAi but not with the control RNAi substantially reduces the level of RhoA protein, supporting the conclusion that the effects of RhoA-directed RNAi on the ability of PC1 to decrease GSK3β kinase activity depicted in Figure 4a is attributable to the suppression of RhoA expression. n≥3 for all experiments. **(b) RNAi-mediated knockdown of G_α13_** Cell lysates were prepared 24 hours after treatment with G_α13_-directed or control RNAi, separated by SDS-PAGE and electrophoretically transferred to a nitrocellulose membrane. The blot was probed with an antibody directed against G_α13_, as well as with an antibody directed against actin, which served as a loading control. Treatment with the G_α13_-directed RNAi but not with the control RNAi substantially reduces the level of G_α13_ protein, supporting the conclusion that the effects of G_α13_-directed RNAi on the ability of PC1 to decrease GSK3β kinase activity depicted in Figure 4a is attributable to the suppression of G_α13_ expression. **(c) G_α13_ is not detected in anti-HA immunoprecipitations from HEK293 cells transfected with empty vector.** Lysates prepared from HEK293 cells transfected with empty vector were subjected to immunoprecipitation with anti-HA antibody and the recovered material was separated by SDS-PAGE, electrophoretically transferred to a nitrocellulose membrane and analyzed by western blotting with antibodies directed against G_α13_ as indicated. G_α13_ is not detected in anti-HA immunoprecipitates from empty-vector transfected HEK293 cells, demonstrating that the co-precipitation of G_α13_ with anti-HA from cells expressing PC1 constructs (Figure 4c and d) is specific. **(d) PC1^ΔGBS^ protein co-expressed with PC2 is present at the surface of HEK293 cells.** Surface immunofluorescence was performed by incubating non-permeabilized HEK293 cells with anti-Flag antibody followed by incubation with an Alexa488-conjugated secondary. Like wild type PC1, the PC1ΔGBS protein co-expressed with PC2 is detectable at the cell surface. **(e) Assessment of expression of PC1 constructs in the samples employed in the fluorescent analysis of GSK3β activity depicted in** Figure 4E. Lysates prepared from HEK293 cells subjected to each of the conditions analyzed in Figure 4f were separated by SDS-PAGE and analyzed by western blotting using antibodies directed against HA to detect the PC1 CTF and against actin as a loading control.

**Supp. Fig. 5A.**
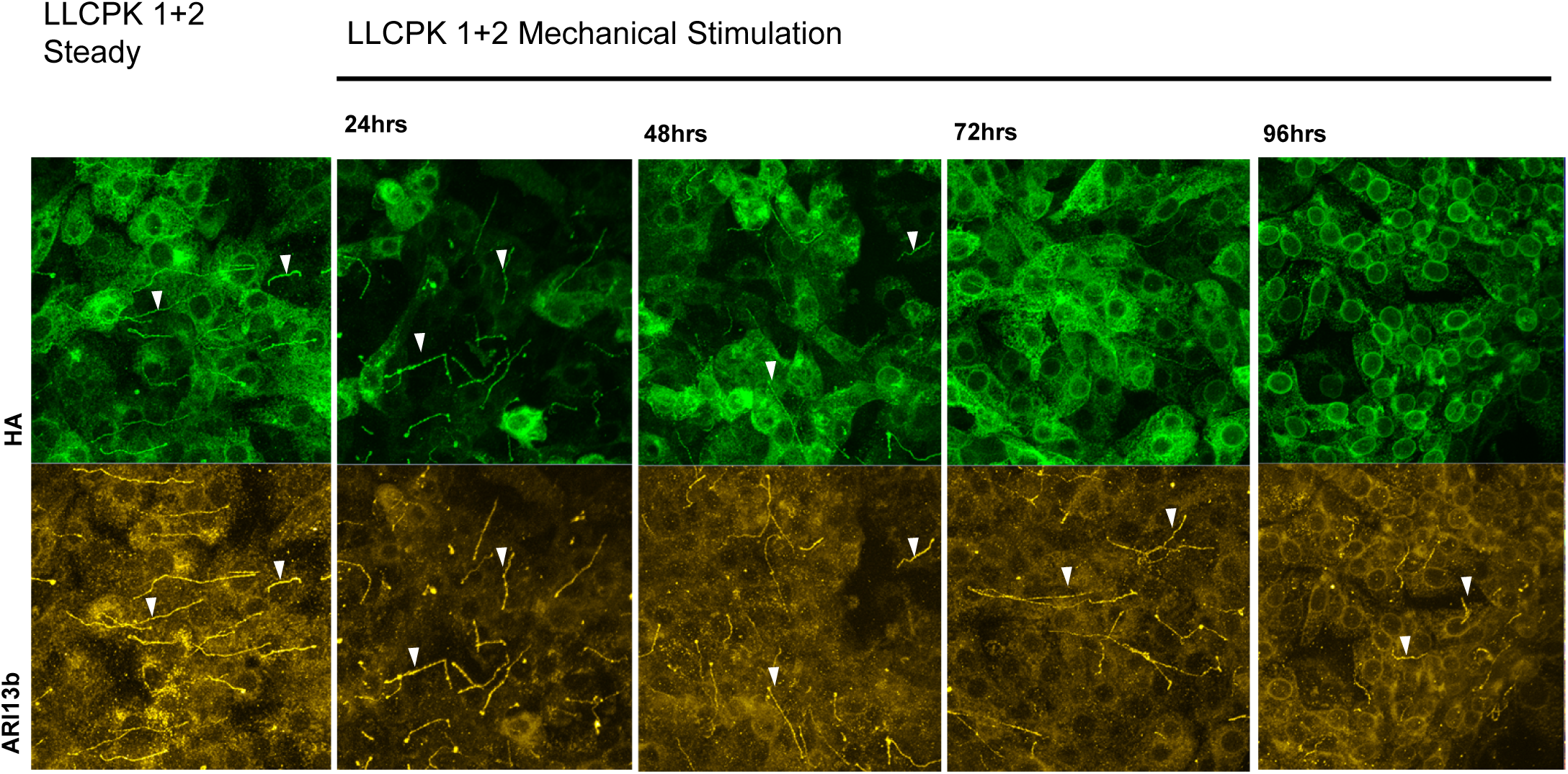
Time course of mechanical stimulus-induced PC1 relocalization from the primary cilia in LLCPK1+2. Prolonged(>24hrs) mechanical stimulation is required to cause PC1 to leave the primary cilia, as demonstrated in mechanically stimulated LLCPK1+2 cell culture.

**Supp. Fig 5B.**
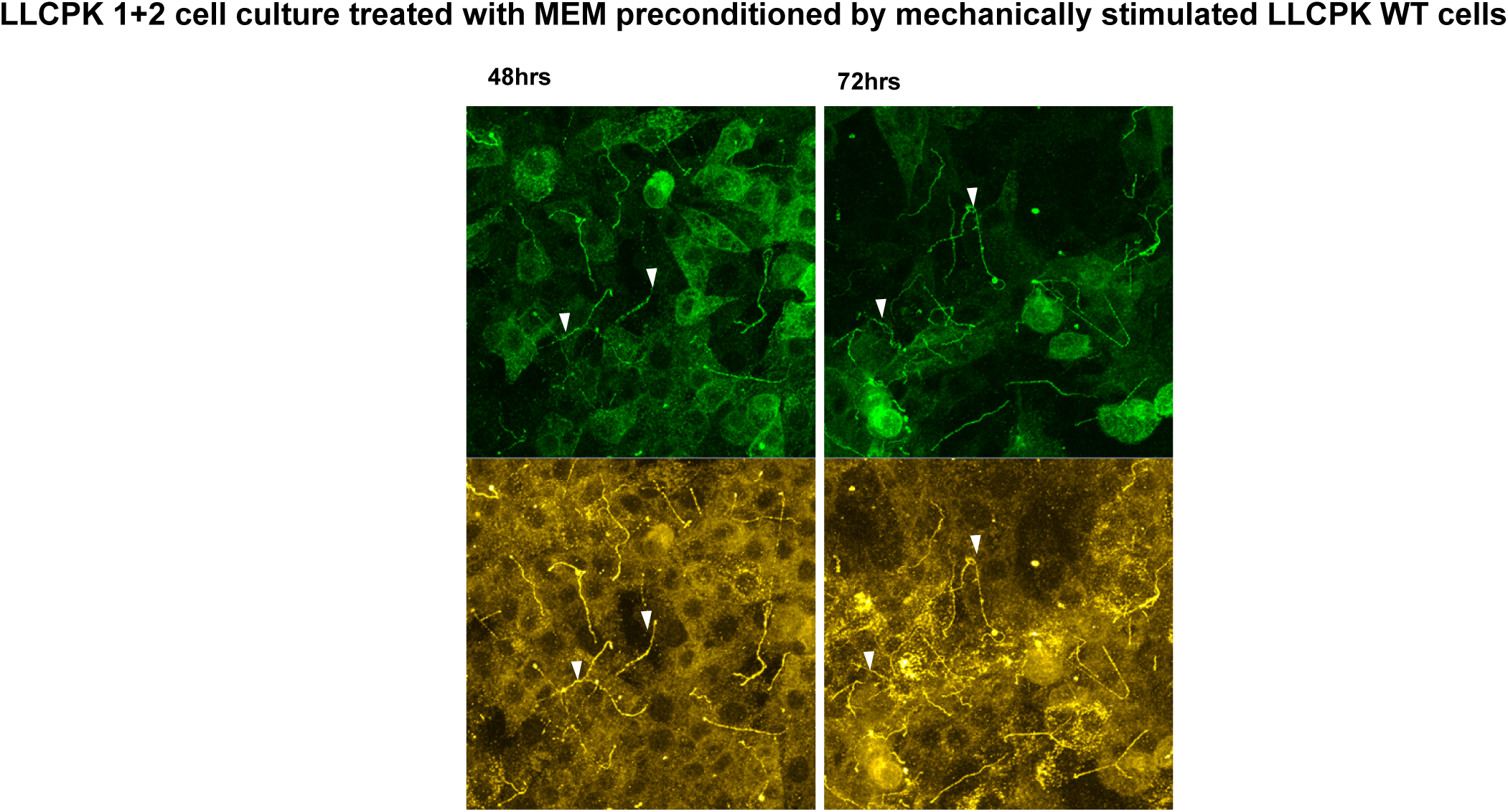
WT LLCPK are unable to promote conditioning of the medium in response to mechanical stimulus. LLCPK1+2 cells were treated with MEM preconditioned by mechanically stimulated LLCPK WT cells. Even after 72hrs of incubation the pool of ciliary PC1 is clearly detectable, demonstrating the lack of secreted signal by WT LLCPK culture subjected to mechanical stimulation. N=3, statistical significance was assessed by paired Student’s T test. *= P<0.05, **= P<0.01, ***=P<0.001

**Supp. Fig. 6.**
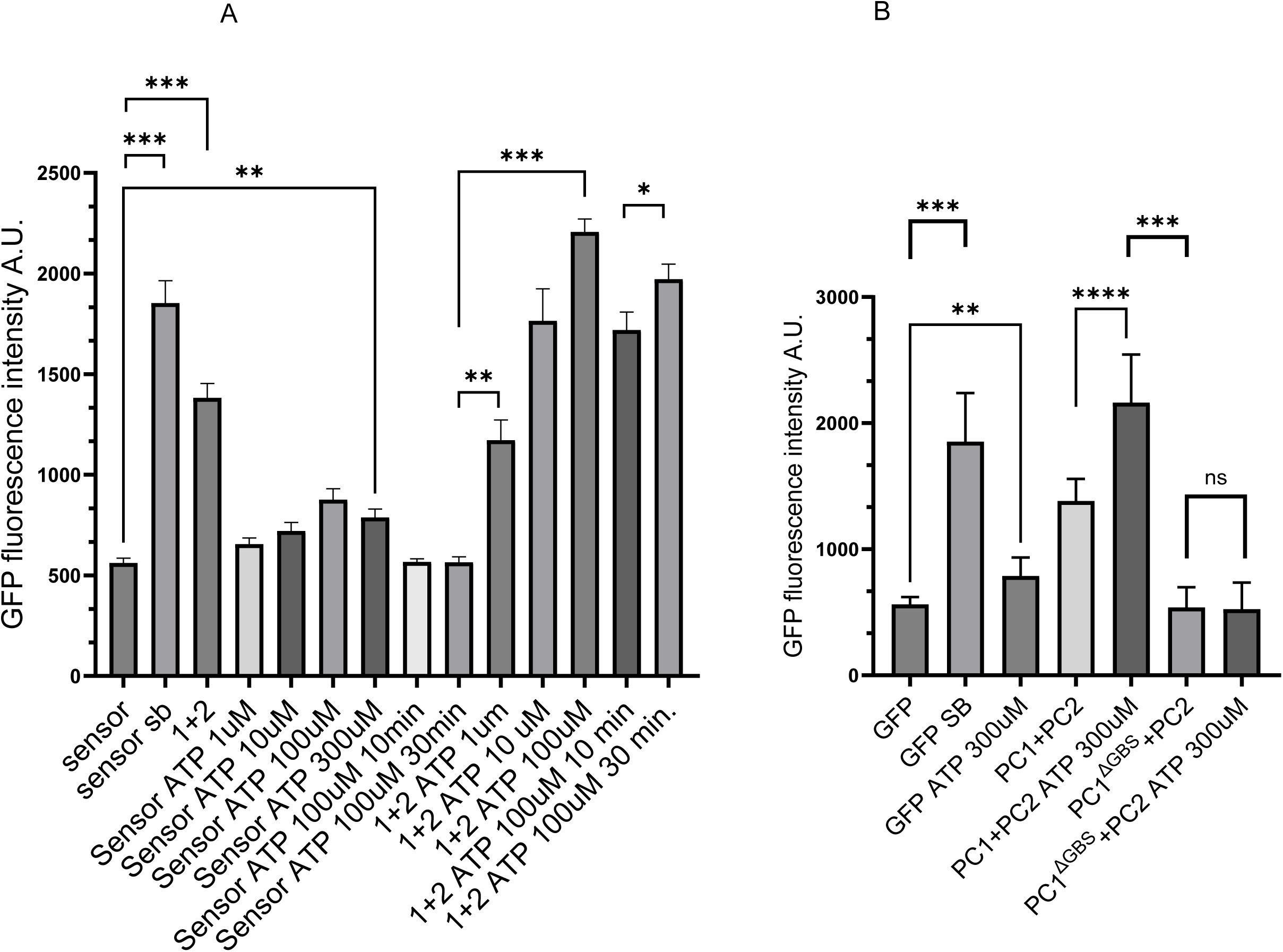
Incubation with ATP leads to GSK3β inhibition through a mechanism that involves the receptor activity of PC1. **(a)** HEK293 cells expressing the GSK3β biosensor with or without PC1 and PC2 were incubated in the absence or presence of the indicated concentrations of ATP for 3 hours or for the indicated times. Half maximal biosensor activation was observed after incubation with 10 µM ATP for 10 minutes. Maximal activation of biosensor signal, exceeding that detected through incubation with the GSK3β inhibitor SB 266763, was observed following incubation with 100 µM ATP overnight. **(b)** To assess the role of the PC1 interaction with trimeric G proteins in the ATP-mediated activation of the biosensor, the assay was performed in HEK293 cells transfected to express wild type PC1 and PC2 or PC1 mutated to lack its G protein binding site (PC1^ΔGBS^) along with PC2. While in cells expressing wild type PC1 and PC2 incubation with 300 µM ATP overnight increased the biosensor signal to the same extent as the GSK3β inhibitor SB 266763, no ATP effect on the biosensor signal was seen in cells that express PC1^ΔGBS^ together with PC2. Thus, PC1 that lacks its G-protein binding site does not inhibit GSK3β upon extracellular stimulation with ATP.

## Notes

### Competing Interest Statement

The authors have declared no competing interest.

